# Antibiotic inhibition uncouples growth from horizontal gene transfer

**DOI:** 10.64898/2026.10.14.759178

**Authors:** Heather D. Curtsinger, Gabriel Quintero, Aidan Larish, Tatiana Dimitriu, Christian Lesterlin, Allison J. Lopatkin

## Abstract

Conjugative plasmids often encode resistance to multiple antibiotics, allowing a single mobile genetic element to transfer across diverse selective environments. This co-selection is often invoked to explain the persistence and spread of multidrug resistance, with antibiotic resistance genes encoded on the same plasmid frequently treated as equivalent routes of plasmid maintenance. However, accumulating evidence suggests that different antibiotics can produce distinct co-selection outcomes. These differences have been interpreted through the lens of fitness and population-level selection, while the potential effects on plasmid transfer itself are largely unknown. Here, we show that different antibiotics generate significantly different transfer dynamics for the exact same multidrug-resistant plasmid. Surprisingly, much of this variation arises from continued conjugation after antibiotic exposure begins, whereby a transient residual conjugation window persists despite strong growth inhibition, allowing new transconjugants to continue forming for hours after treatment in an antibiotic dependent manner. A simple model for describing antibiotic effect on transfer competence of both donors and recipients quantitatively explains these antibiotic-specific dissemination dynamics. Moreover, different recipient genetic backgrounds exhibit distinct residual conjugation windows, demonstrating that the capacity for continued plasmid transfer during antibiotic exposure is both drug- and host-dependent. Together, our results show that antibiotics suppress plasmid dissemination on different timescales, creating drug-specific residual conjugation windows that are largely decoupled from growth inhibition. These findings identify conjugation dynamics as a previously overlooked contributor to co-selection outcomes and suggest that antibiotics that select for the same multidrug-resistant plasmid are not equivalent in their ability to promote plasmid persistence.

## Introduction

Horizontal gene transfer (HGT) enables bacteria to rapidly acquire new genetic material from nearby cells, accelerating microbial adaptation beyond what mutation and clonal expansion alone can achieve(*1*). Of the many types of HGT, plasmid conjugation, which describes the transfer of plasmids via direct cell-to-cell contact, plays a central role in the evolution and dissemination of antimicrobial resistance genes (ARGs)(*2–6*). Understanding the factors that govern conjugation is critical for predicting and controlling the spread of resistance in clinical and environmental settings.

Among the many environmental factors that influence the dissemination of plasmids that encode ARGs(*7–11*), antibiotics play a uniquely important role since they also determine the selective advantage of plasmid carriage. This influence is particularly pronounced for multidrug-resistant plasmids, which can encode resistance to multiple antibiotics simultaneously and therefore can persist across diverse selective environments(*12*). Indeed, exposure to any one antibiotic can enrich for the whole plasmid, facilitating the maintenance of the full resistance cassette it carries(*13*, *14*). This phenomenon, commonly referred to as co-selection, represents a central framework for understanding the persistence and spread of multidrug-resistant plasmids. For example, co-selection is routinely invoked to explain why resistance determinants persist despite reduced use of individual antibiotics(*15*), why stewardship efforts do not always produce corresponding declines in resistance prevalence(*16*), and how exposure to a single antibiotic can maintain an entire multidrug-resistant gene cassette through genetic linkage(*15*, *17*, *18*).

In the context of co-selection, different antibiotics are often assumed to represent equivalent routes of plasmid maintenance(*19*, *20*). However, evidence suggests that different antibiotics do not necessarily select multidrug-resistance elements equivalently(*21*, *22*). For example, one study showed that the same multidrug-resistant plasmid could be selected across different antibiotics and heavy metals at varying levels of magnitudes(*14*). Similarly, another study showed that tetracycline selected for the same multidrug-resistant plasmid at concentrations nearly 100-fold lower relative to the MIC than ampicillin, a difference attributed to the population-level effects of their respective resistance mechanisms(*23*). Community-level studies have likewise reported preferential enrichment of specific resistance determinants despite extensive genetic linkage among co-occurring ARGs(*24*). Together, these observations suggest that antibiotics may differ substantially in their ability to maintain and enrich multidrug-resistant elements.

The basis for these asymmetric co-selection outcomes has largely been interpreted through the lens of fitness and population-level selection dynamics. However, in addition to being maintained within their host, the persistence of multidrug-resistant plasmids also depends on their continued spread into new cells through conjugation. Consequently, antibiotic-specific differences in plasmid conjugation rates themselves may represent an underappreciated contributor to co-selection dynamics. Indeed, previous studies have shown that antibiotics can influence conjugation transfer rates through a variety of mechanisms, including direct regulation of transfer functions and indirect effects on donor and recipient populations(*25–27*). However, these studies have largely examined the effects of individual antibiotics on a given plasmid, revealing whether a particular drug increases or decreases plasmid transfer relative to a no-drug control. Whether different antibiotics selecting for the same multidrug-resistant plasmid exert equivalent effects on plasmid transfer is largely unknown. Resolving this can reveal how antibiotic identity shapes the persistence and spread of multidrug resistance.

Here, we show that different antibiotics, to which the recipient is sensitive, that select for the same multidrug-resistant plasmid, significantly alter the amount of transconjugants that form. Surprisingly, much of this variation arises from conjugation that persists after antibiotic exposure begins, despite strong inhibition of bacterial growth. This residual conjugation window occurs with both bacteriostatic and bactericidal antibiotics and across multiple plasmid types. A simple quantitative model in which donor and recipient populations contain transient but declining pools of transfer-viable cells quantitatively explains these antibiotic-specific effects. Finally, we show that different recipient strains also exhibit unique residual conjugation capacities, revealing a previously unrecognized strain-specific advantage for plasmid dissemination under drug selection. Together, our results demonstrate that antibiotics inhibit growth more rapidly than they inhibit HGT, creating drug-specific residual transfer windows that cause the same multidrug-resistant plasmid to spread differently depending on the antibiotic and its concentration. These findings identify residual conjugation dynamics as a previously overlooked factor shaping plasmid transfer under antibiotic selection, with implications for how different antibiotics may differentially co-select the same multidrug-resistant plasmid.

## Results

### Antibiotic identity impacts conjugation efficiency for the same multidrug resistance plasmid

Different antibiotics can impose distinct selective outcomes on the same multidrug-resistant plasmid. However, whether such differences can arise due to drug-specific effects on the transfer of that plasmid, separate from the growth of transconjugants, is unknown. Addressing this is challenging since conjugation assays typically combine plasmid transfer, antibiotic selection, and transconjugant quantification in a single measurement. Thus, differences in conjugation rates may reflect differences in transfer, survival under selection, or both. To distinguish these possibilities, we used a matched setup consisting of “mixture” and “background” conditions (**Fig. 1A**). In the “mixture”, donors and recipients were first incubated together for one hour in a non-selective environment before selective plating, allowing transconjugants to form prior to antibiotic exposure. In the parallel “background”, donors and recipients were incubated separately under the same conditions and mixed only at the time of plating; any transconjugants from the background thus arise from transfer occurring on the agar surface after antibiotic exposure began (See **Methods** for more details). Comparing these matched conditions allows us to isolate antibiotic-specific effects on plasmid transfer occurring under selection.

**Figure 1.**
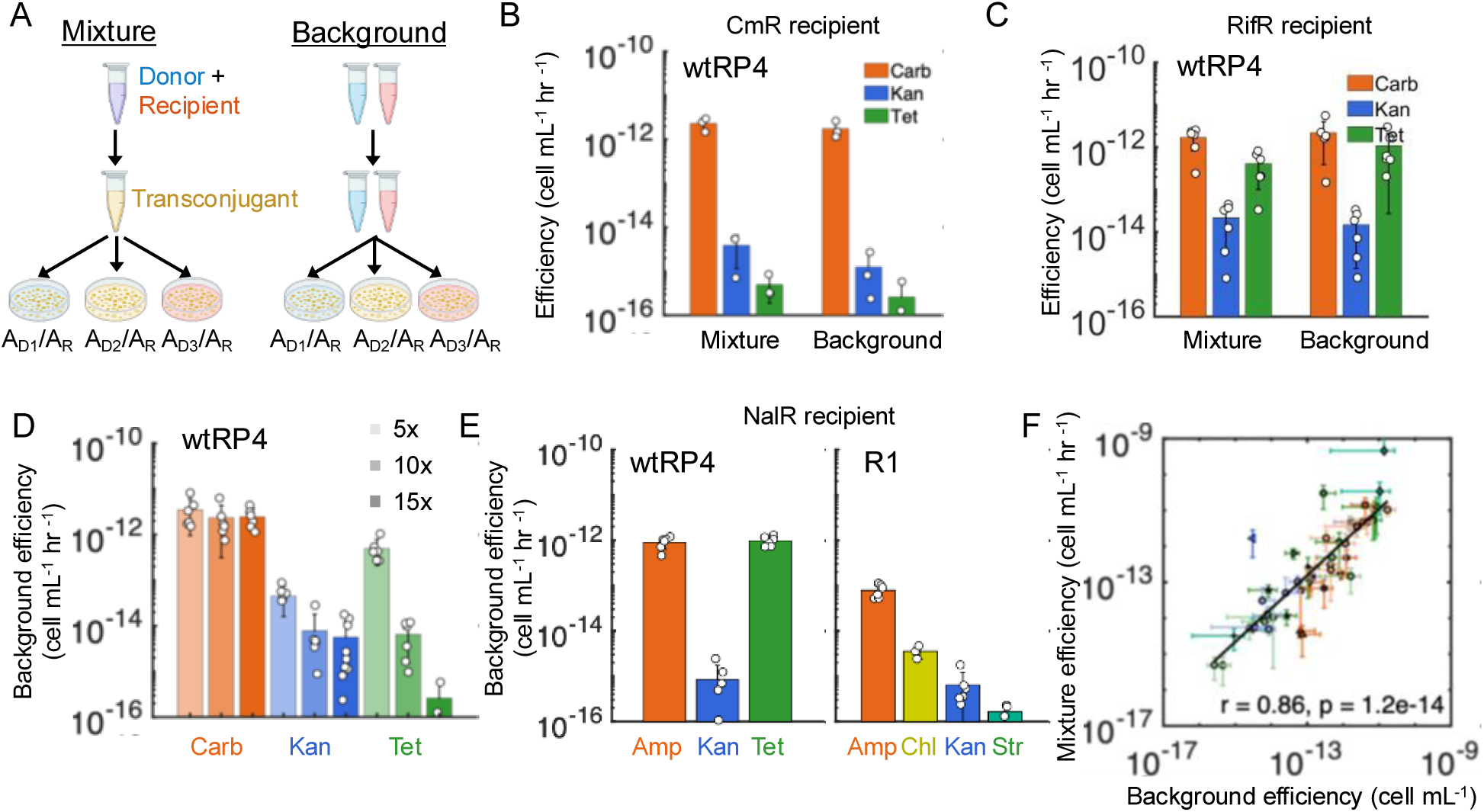
Antibiotic selection changes the observed conjugation efficiency. **A** Schematic of the experimental set-up for mixture and background conditions. Donor and recipient cells were either mixed together (mixture) or kept separate (background) during incubation (1 hour at 25°C). Following incubation, both conditions are diluted and plated onto agar plates containing dual antibiotics for selection of transconjugants. A_D1_, A_D2,_ and A_D3,_ represent 3 different drugs used to select for transconjugants of the same donor plasmid. A_R_ represents the antibiotic that the recipient encodes resistance for. **B** Conjugation efficiency under different antibiotic selection conditions using BWcmR recipients and BWrifR-wtRP4 donors. Cells were plated onto agar containing 5x Cm mixed with 15x Carb (orange), Kan (blue), or Tet (green). Conjugation efficiency is plotted on the y-axis (see Methods) for the mixture (*left*) or background (*right*) condition. The bar height represents the average of at least three biological replicates (scatter points). Drugs significantly impact the mixture (one-way ANOVA, P = 2.5×10^-5^) and background (P = 5.5×10^-5^). There’s no difference between the mixture and background for Carb (two-tailed paired t-test with Bonferroni correction, P = 1), Tet (P = 0.3), and Kan (P = 0.2) (n = 3). **C** Conjugation efficiency under different antibiotic selection conditions using BWrifR recipients and DA-wtRP4 donors. Cells were plated onto 5x Rif mixed with 15x Carb (orange), Kan (blue), or Tet (green). Conjugation efficiency is plotted on the y-axis (see Methods) for the mixture condition (*left*) or in the background condition (*right*). Drugs significantly impact the mixture (one-way ANOVA, P = 3.1×10^-5^) and background (P = 3.7×10^-6^). There’s no difference between the mixture and background for Carb (two-tailed paired t-test with Bonferroni correction, P = 1), Tet (P = 0.4), and Kan (P = 0.06) (n = 6). **D**. Effect of antibiotic selection dose on background conjugation efficiency. The background condition for recipient BWcmR and donor BWrifR-wtRP4 was plated on 15x, 10x, or 5x the MIC of the plasmid selecting antibiotic (Carb, Kan or Tet). The doses had no significant association with background efficiency for Carb (Spearman, ρ = -0.16, P = 0.5, n = 18) but was significant for Tet (ρ = -0.9, P = 4.5×10^-7^, n = 15) and Kan (ρ = -0.5, P = 0.02, n = 21). **E** Background conjugation efficiency across antibiotic selection conditions using NalR recipients. The background efficiency was calculated from previously published(*22*) data using wtRP4 (*left*) and R1 (*right*) with a nalidixic acid resistant (NalR) recipient under different conjugation conditions (3 hours, shaking, 37°C). Antibiotics significantly impact the background for wtRP4 (one-way ANOVA, P = 3.2×10^-11^) and R1 (P = 2.4×10^-11^). **F** Relationship between the mixture and background conjugation efficiencies across plasmids and conjugation conditions. Mixture efficiency (y-axis) is plotted against background efficiency (x-axis) across all different plasmids, conjugation conditions, donors, and recipients (n = 46). A strong positive correlation was observed (Pearson, r = 0.86, P = 1.2×10^-14^).

To enable direct comparison across antibiotics, a single donor-recipient mixture (or corresponding individual parents) was distributed onto different selective plates, requiring a plasmid capable of generating transconjugants both in liquid (during the incubation period) and on solid (after plating) media. We therefore selected the multidrug-resistant incompatibility group P (IncP) plasmid RP4, which encodes resistance to kanamycin (Kan), carbenicillin (Carb), and tetracycline (Tet), and conjugates efficiently on solid surfaces(*22*, *28*). To confirm that transconjugants were already present before plating (e.g., after static liquid incubation), we used a fluorescent RP4 derivative (sfRP4) that activates GFP expression only after transfer into the *Escherichia coli* recipient strain BW25113 with rifampicin (Rif) resistance (referred herein as BWrifR), but remains repressed when carried by an *E. coli* donor expressing constitutive TetR (referred to as MG-sfRP4). After verifying that pure BWrifR-sfRP4 transconjugants with constitutive GFP fluorescence could be distinguished from non-fluorescent recipients and GFP-repressed donors (**Fig. S1A**), we performed a standard conjugation reaction (minimal media and static incubation for 1 hour at 25°C(*29*)). The mixture’s fluorescence was then quantified via flow cytometry. Newly-formed GFP-positive cells were readily detected in the mixture (**Fig. S1B**), confirming that transconjugants were present in the mixture prior to antibiotic selection.

We next used the wild-type RP4 plasmid (wtRP4) carried by BWrifR (e.g., BWrifR-wtRP4 donors), and the *E. coli* strain BW25113 carrying a small plasmid pZE31-GFP, encoding for constitutive green fluorescent protein (GFP) and chloramphenicol (Cm) resistance, referred herein as BWcmR) as the recipient, for assessing the effects of different antibiotics on RP4 conjugation. Antibiotic concentrations were first standardized relative to the minimum inhibitory concentration (MIC) of the corresponding sensitive parent strain (**Table S1**). As an initial concentration choice, Cm was used at 5x the donor MIC for recipient (BWcmR) selection, and all three antibiotics were used at 15x the recipient MIC for donor (BWrifR-wtRP4) (**Fig. S2A, S2D**). Relatively high concentrations for plasmid selection were initially used because we reasoned that strong inhibition would rapidly suppress bacterial growth and limit the opportunity for antibiotic-specific effects on conjugation to emerge, whereas lower Cm concentrations were sufficient for donor counterselection and minimized recipient-specific growth effects unrelated to conjugation. We quantified the conjugation efficiency under each drug condition using the standard bimolecular reaction equation (see **Methods,** “*Conjugation Assay” section*), thus accounting for any discrepancies in donor/recipient densities(*30*).

BWrifR-wtRP4 conjugation efficiencies in the mixture differed significantly depending on the antibiotic used for selection (**Fig. 1B**, *left*, one-way ANOVA, P = 2.6×10^-5^). The highest efficiency was observed with Carb, followed by nearly 2-log and a further 1-log reduction with Kan and Tet, respectively. Moreover, mixture conjugation efficiencies for all three antibiotics closely tracked the corresponding background efficiency (**Fig. 1B**, *right*, one-way ANOVA, P = 5.5×10^-5^), with no statistical difference between any paired drug mixture and background efficiencies (two-tailed paired t-test with Bonferroni correction Tet: P = 0.3, Carb: P = 1, Kan: P = 0.2, n = 3); this indicates that the majority of observed transconjugants were formed under selective plating instead of during the 1 hour mating window. While wtRP4 conjugation on solid agar is itself not surprising(*22*, *28*, *31*), each antibiotic selection within a single condition (e.g., mixture or background) began with the same donor-recipient mixture. Importantly, the occurrence of colonies in the background required the presence of both mating partners, as donor-only and recipient-only controls produced no colonies under these conditions (**Fig. S3A**). Thus, the observed antibiotic-specific differences in conjugation efficiency are unlikely to arise solely from differential growth of pre-existing transconjugants and instead suggest differences in conjugation events during drug exposure.

### Antibiotic-dependent background conjugation is not explained by differences in viable cell counts or plating artifacts

Alternative sources could also contribute to the observed background signal, including differences in cell survival or technical effects introduced by the plating protocol. Thus, we first confirmed that neither the BWrifR-wtRP4 donor strain nor the established wtRP4 transconjugants (BWcmR-wtRP4) exhibited antibiotic-specific effects on selective plates (**Fig. S3B**, both P > 0.9, one-way ANOVA), ruling out differential plate effects as the source of the observed differences in conjugation efficiency. Next, we confirmed that colonies formed under background conditions were not spontaneous donor mutants (**Fig. S3C**); together with the absence of colonies in recipient-only controls (e.g., **Fig. S3A**), this excludes spontaneous resistance in either parental population as the source of the signal. Finally, reversing the order of donor and recipient spotting had no effect on transconjugant yield (**Fig. S3D**), and replacing spot plating with spread plating, which eliminates any prolonged residence of cells in a liquid droplet, produced similar patterns in background efficiencies across drugs (**Fig. S3E**), indicating that plating method does not bias the drug-specific outcome. Together, these findings confirm that differences in background conjugation arise from antibiotic-specific effects after plating under our experimental conditions.

### Background transconjugant formation is robust across strains, plasmids, and conjugation conditions

To determine if these antibiotic-specific differences were unique to Cm selection, we first repeated the assay using two additional recipient strains, BWgentR and BWrifR, carrying resistance markers for gentamicin (Gent) and rifampicin (Rif), respectively (see **Table S2** for all strains); to enable the use of Rif recipient selection, the donor DA-wtRP4 was used instead, a closely related strain to BW25113 without Rif resistance. Using the same MIC-fold selection levels as above, we again observed that the choice of antibiotic used to select for the plasmid led to distinct conjugation efficiencies (one-way ANOVA, BWgentR mixture: P = 5×10^-5^, background: P = 7.5×10^-5^ and BWrifR mixture: P = 3.1×10^-5^, background: P = 3.7×10^-6^) (BWrifR: **Fig. 1C**, BWgentR: **Fig. S4A**); moreover, as before, these trends were largely consistent between mixture and background measurements. Thus, differences in background conjugation efficiencies across plasmid antibiotic selection are not specific to the recipient drug used for selection.

Although a concentration of 15x the MIC is far above levels required for growth inhibition, the drug concentration may still affect antibiotic efficacy. Indeed, using the Cm-resistant recipient BWcmR, we tested varied plasmid-selecting antibiotic concentrations at 5x, 10x, and 15x MIC, and found that conjugation efficiency decreased with increasing concentrations for Kan (Spearman, ρ = -0.5, P = 0.02, n = 21) and Tet (ρ = -0.9, P = 4.5×10^-7^), whereas no significant concentration dependence was observed for Carb (**Fig. 1D**;= ρ = -0.16, P = 0.5). Likewise, decreasing the recipient antibiotic concentration (5x, 4x, 3x, and 2x MIC of Cm), while holding plasmid selection constant (15x), increased the conjugation efficiency by ∼1-log (**Fig. S4B**; Spearman ρ = -0.7, P = 0.007). Finally, dose dependence was not unique to Cm, as it was also observed using Tet/Rif system, even with higher Rif (6x donor MIC) (**Fig. S4C**). Thus, background conjugation occurs in a largely dose-dependent manner, although the magnitude of this change depends on the specific antibiotic.

To determine if these findings were dependent on the physiology of our assay conditions, we leveraged recently published data that also examined similar background setups but using a different protocol whereby cells were incubated shaking for 3 hours in rich media at 37°C prior to selection with the indicated donor drug and nalidixic acid for recipient selection(*22*). Despite these differences, for wtRP4, Ampicillin (a β-lactam antibiotic closely related to Carb) and Tet still yielded higher efficiencies than Kan (∼10^-12^ cells/ml/hr vs ∼10^-15^ cells/mL/hr) (**Fig. 1E**, *left*; one-way ANOVA, P = 3.2×10^-11^). Moreover, a second conjugative plasmid R1, belonging to a different incompatibility group (IncFII), also exhibited antibiotic-dependent differences in conjugation efficiency under these conditions (**Fig. 1E**, *right;* one-way ANOVA, P = 2.3×10^-11^), indicating these results are not specific to the IncP plasmid type.

Finally, to examine the generality of this observation, we compared mixture and background conjugation efficiencies across all measurements, including 14 plasmids, 2 assay methods, 5 recipients, 5 antibiotics, multiple concentrations, and different media types (46 total conditions; **Table S3**), revealing the two values remained strongly correlated (Pearson, r = 0.86, P = 1.2×10^-14^) (**Fig. 1F**). Despite this consistent relationship, antibiotic identity produced nearly five orders of magnitude variation in conjugation efficiency depending on the specific condition. Together, these results demonstrate that antibiotic-specific differences in conjugation arise largely from differences in continued plasmid transfer after antibiotic exposure begins.

### Plasmid transfer persists under growth-inhibitory conditions

The observed antibiotic-specific differences in continued plasmid transfer could arise primarily during the initial period before antibiotics exert their full inhibitory effects, or persist after population growth has largely stopped. To distinguish these possibilities, we measured background conjugation after progressively longer periods of antibiotic exposure of the donors and recipients individually (**Fig. 2A**). This directly quantifies the duration over which parental cells remain conjugation-competent following antibiotic treatment.

**Figure 2.**
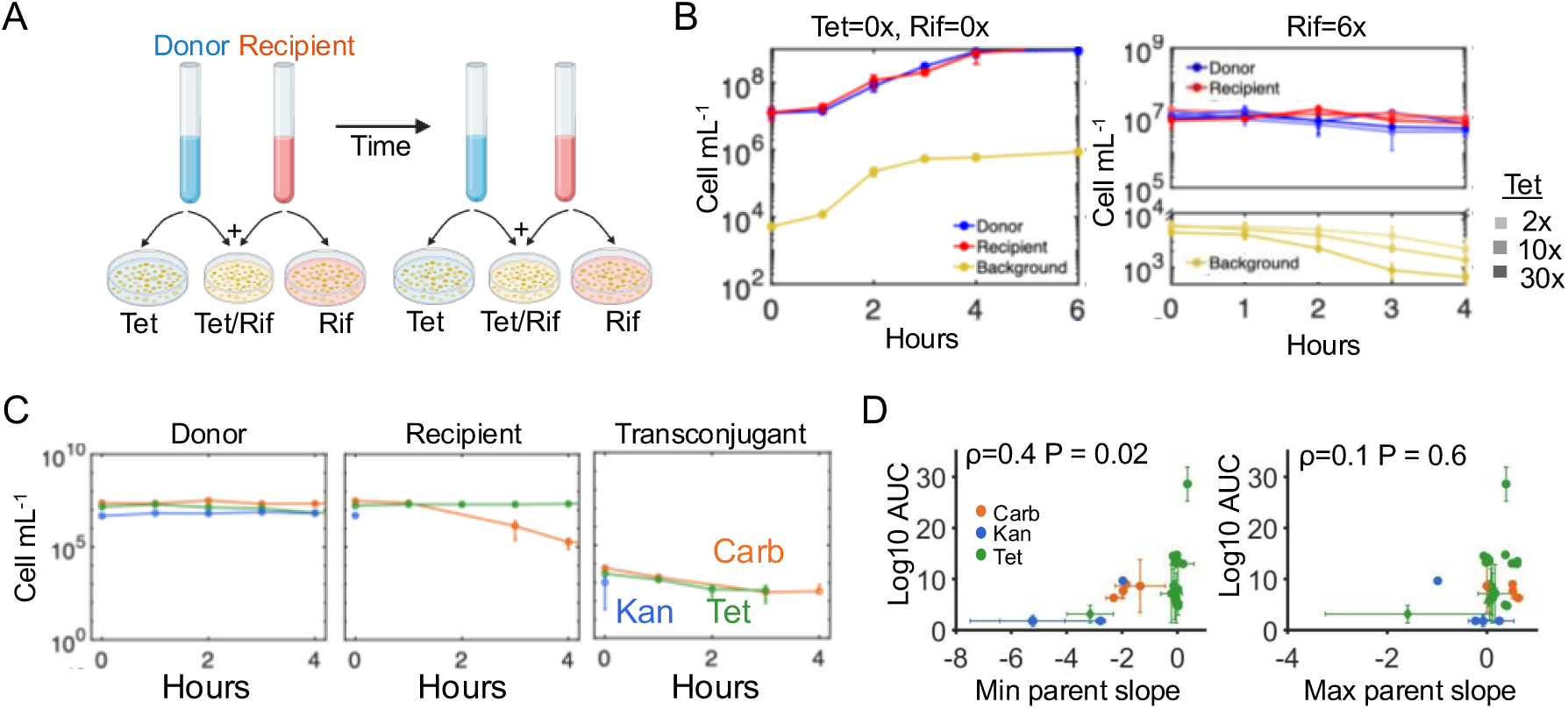
Plasmid transfer persists under inhibitory antibiotic concentrations. **A** Schematic of background time courses. Donors and recipients were statically incubated for 1 hour at 25°, followed by separately diluting into tubes containing LB media and indicated antibiotic. Aliquots are removed at various time points (0-6 hours), and donors and recipients are spotted on top of each other onto dual antibiotic agar for transconjugant enumeration, and individually for parent density quantification. **B** Background time courses under no antibiotics (*left*) and increasing tetracycline exposure (*right*). Transconjugants (yellow), donor DA-wtRP4 (blue), and recipient BWrifR (red) cell counts per mL are plotted on the y-axis for both conditions. The background time course under no antibiotic was collected at 0, 1, 2, 3, 4, and 6 hours (*left*), and with antibiotic at 0, 1, 2, 3, and 4 hours (*right*). For the antibiotic condition, Rif is used at 6x MIC and increasing doses of Tet at 2x, 10x, or 30x MIC (dark to light shading). Increasing Tet doses are negatively correlated with cell counts per mL (Spearman, ρ = -0.79, P = 0.002, n = 3). **C** Background time courses under different donor-selecting antibiotics. Background time courses were collected as described in (B) using Donor DA-wtRP4 and recipient BWrifR. Antibiotic conditions include 6x Rif with either 15x of Carb (orange), Tet (green), or Kan (blue). In all cases, transconjugants were plated onto Tet/Rif to isolate tube-specific effects. The background time course for the donor individually (*left*), recipient (*middle*), and transconjugants (*right*) are shown under these conditions (n = 3, for each panel). **D** Relationship between transconjugant growth and parental growth from time course curves. The area under the curve (AUC) was calculated from all collected background curves (y-axis) and plotted against the minimum (*left,* Spearman, ρ = 0.4, P = 0.02) or maximum (*right*, ρ = 0.1, P = 0.6) slope of the parents. The shape of the marker represents the recipient antibiotic used (circle for Rif, triangle for Cm, and diamond for Kan selection) and the color represents the donor antibiotic used (blue for Kan, green for Tet, and orange for Carb). For example, pB10 is indicated by a green diamond. The full list containing all the conditions plotted can be found in **Table S4** (n = 28).

First, we used the DA-wtRP4 donor and the BWrifR recipient as above (e.g., **Fig. 1C**) and confirmed that background conjugation remained detectable over time when neither parent was inhibited by an antibiotic. Under these conditions, both donor and recipient populations grew in liquid culture, as expected (**Fig. 2B**, *left*, red and blue lines). Consistent with this, background conjugation was observed at every time point, with transconjugants monotonically increasing (**Fig. 2B**, *left*, yellow line).

We next treated the donors and recipients each with both Tet and Rif inhibition (Tet at doses 2x, 10x, 30x MIC and Rif at 6x MIC). Both parents remained approximately constant over time, consistent with static antibiotic action (**Fig. 2B**, *right*). Despite the absence of net population growth, transconjugant colonies continued to form for up to 4 hours, demonstrating that conjugation persisted well beyond growth arrest. This trend was dose-dependent, as the amount of background conjugation reduced with increasing Tet concentrations in the tube (**Fig. 2B**, *right*; Spearman ρ = -0.79, P = 0.002). Importantly, this concentration dependence mirrored that observed in the original single-time-point measurements (e.g., **Fig. S4D**), suggesting that differences captured by the single background measurement may reflect variations in the magnitude and/or duration of post-drug residual conjugation.

We therefore asked whether the antibiotic-specific differences observed in the single-time-point assay similarly corresponded to distinct residual conjugation dynamics. We quantified conjugation over time following exposure of donors and recipients to 15x Kan, Tet, or Carb together with 6x Rif, as above. All transconjugants were plated onto the same Tet/Rif plates so that differences in transconjugant recovery could be attributed to antibiotic exposure in the parental populations during liquid culture rather than to plate selection. Consistent with the rank ordering observed in the original single-time-point measurements (**Fig. 1C**), Carb and Tet produced the largest and longest residual windows, while Kan produced the smallest, with recipients falling below our detection limit of 10 cells after 1 hour (**Fig. 2C**). Thus, antibiotic-specific differences captured by a single background measurement seem to reflect distinct residual conjugation windows that persist following antibiotic-mediated growth inhibition.

To determine whether differences in residual conjugation could be explained by parent growth or inhibition alone, we examined 28 combinations of wtRP4-carrying donors and compatible recipients (**Table S4**). For each condition, we estimated donor and recipient growth rates over the residual conjugation window and compared their minimum and maximum slope values with the area under the residual conjugation curve (**Fig. 2D**). Under bactericidal conditions (Carb and Kan), greater parent killing was associated with reduced residual conjugation, resulting in a significant correlation with the minimum parent growth rate (Spearman r = 0.8, P = 0.03), but not the maximum (r = -0.01, P = 1). In contrast, under bacteriostatic conditions (Tet), residual conjugation varied substantially despite near-zero parent growth, with no significant correlation with either growth measure (minimum: r = 0.09, P = 0.7; maximum: r = -0.03, P = 0.9). Across all conditions, residual conjugation correlated weakly with the minimum parent growth rate (r = 0.4, P = 0.02), but not the maximum (r = 0.1, P = 0.6). Together, these results demonstrate that although bacterial killing limits residual conjugation, growth inhibition alone is insufficient to predict its magnitude, revealing a fundamental disconnect between the inhibition of bacterial growth and the inhibition of plasmid transfer.

### A simple residual conjugation window model explains post-antibiotic conjugation dynamics

Continued conjugation after antibiotic treatment suggests that antibiotic-mediated growth inhibition and conjugation inhibition are at least partially separate processes. Moreover, the distinct antibiotic-specific estimates of conjugation efficiency for the same plasmid, even at equivalent MICs, further imply that this drug effect is specific to each antibiotic and not an intrinsic property of the plasmid being transferred. Combined, we hypothesized that separate from growth inhibition, post-antibiotic conjugation is driven by a sub-population of donor and recipient populations that each remains conjugation-competent and declines over time in a drug-dependent manner.

To test this idea, we developed a simple mathematical model (**Fig. 3A**) in which the Donor (D) and Recipient (R) populations are each inhibited by their respective alternative antibiotics (e.g., D is sensitive to A_R_ and R is sensitive to A_D_). Further, we assume that antibiotic treatment results in a decaying viable “conjugation competent” fraction of D and R, denoted F_D_ and F_R_ and parametrized by the first order rate constants k_D_ and k_R_ (**Fig. 3B**). Transconjugants (T) are generated by the interaction between conjugation-competent donors (D_C_=F_D_D) and recipients (R_C_=F_R_R), according to classic bimolecular conjugation kinetics with rate constant η, known as the conjugation efficiency; that is, T= ηR_C_D_C_. Therefore, as *F_R_*, *F_D_*, or both decline, the production of new transconjugants also declines. Moreover, donor, recipient, and transconjugant populations can grow or decline according to their antibiotic-dependent growth rates. Together, these dynamics are described by a five-equation model (**Eqs. 1-5**).

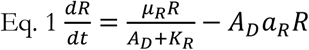

**Figure 3.**
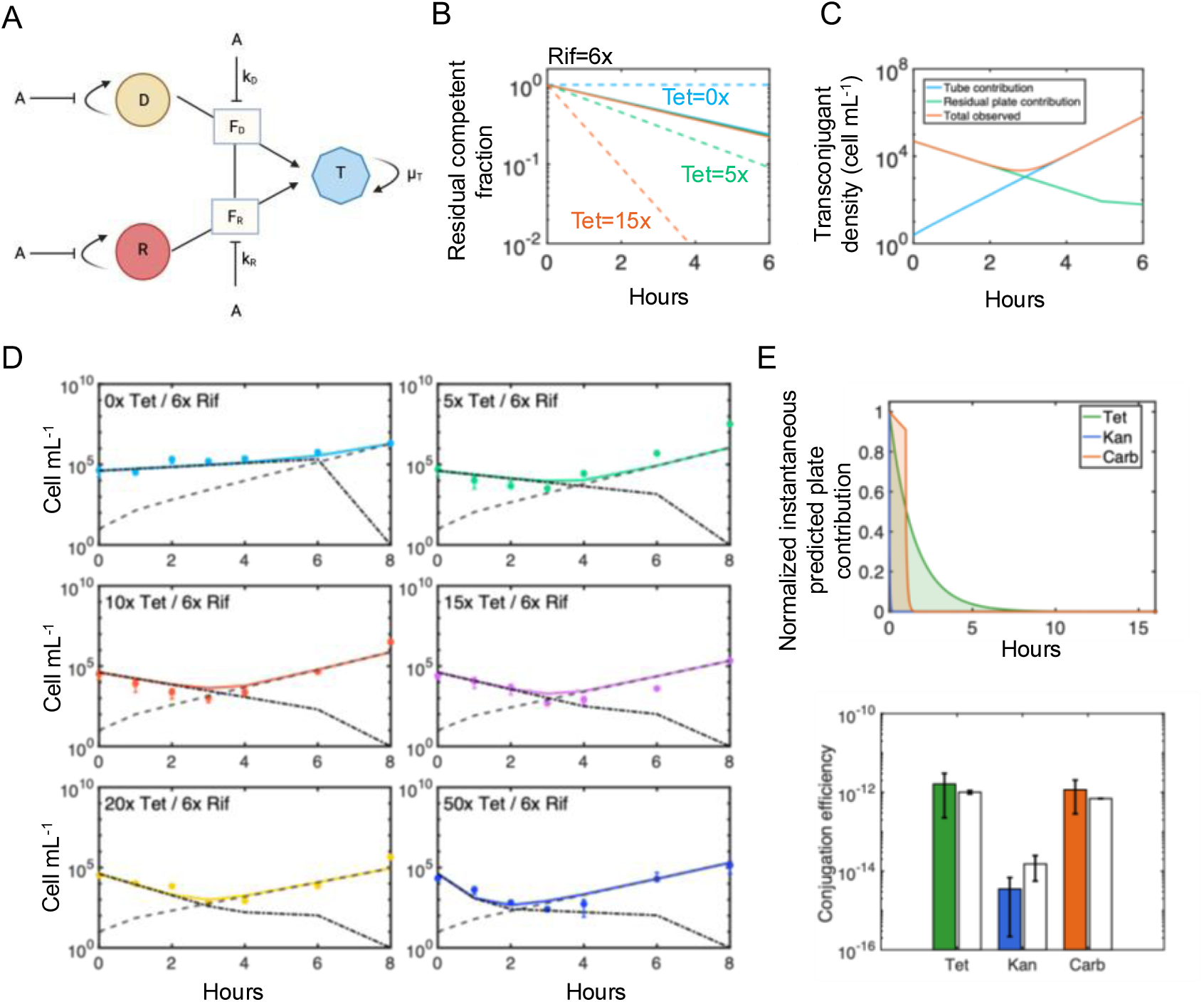
A simple residual conjugation model explains the observed dynamics. **A** Schematic of the model framework. Donors (D) and recipients (R) contain transient transfer-competent fractions F_D_ and F_R_ that decay under antibiotic exposure, generating transconjugants (T) during a finite residual conjugation window. **B** Simulated decay of transfer-competent recipient fractions under increasing tetracycline exposure; 0x Tet (blue), 5x Tet (green), 15x Tet (orange). **C** A representative decomposition of modeled transconjugant dynamics into residual plate contribution (green), tube contribution (blue), and total observed transconjugants (orange). **D** Time courses for the mixtures of Donor DA-wtRP4 and recipient BWrifR. Briefly, donor and recipient were mixed together and incubated for 1 hour at 25°C to allow conjugation to occur and then immediately diluted 75x into 6mLs of LB containing 6x rifampicin and 0x, 5x, 10x, 30x, and 50x tetracycline. Aliquots were removed at time points (0, 1, 2, 3, 4, 6, and 8 hours) and plated onto agar containing 6x Rif and 15x Tet. Data is shown as markers, the dotted line is the predicted plate contribution, the dashed line is the predicted tube contribution, and the solid-colored line is the total predicted Transconjugants (plate+tube). Experiments were conducted in biological triplicate. **E** Predicted time-dependent contribution to conjugation efficiency for Tet/Rif, Kan/Rif, and Carb/Rif plate conditions (*top*). Curves show the model’s instantaneous contribution to cumulative conjugation efficiency, normalized to its value at plating (t = 0), such that each condition begins at 1. Shaded regions indicate the area below each normalized trajectory. Declines reflect the modeled effects of antibiotic-dependent recipient survival and loss of conjugation competence over the plate incubation period. Comparison of predicted and measured conjugation efficiencies across antibiotic conditions recapitulates the data (*bottom*).

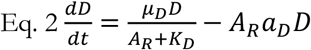

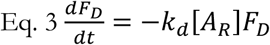

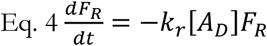

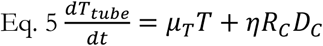

We used Eqs. 1-5 to simulate conjugation during antibiotic exposure in liquid. At each experimental sampling time, we then calculated the additional conjugation occurring after plating by integrating the residual conjugation reaction over the plate incubation period (**Eq. 6; Methods**); here, we assume that the antibiotic-dependent rate of viability loss is the same, but the conjugation efficiency is 100x higher on the plate, based on extensive literature for the RP4 plasmid(*22*, *28*, *32–34*). The total predicted transconjugant population at each time point was therefore the sum of transconjugants generated before and after plating (*T=T_tube_+T_plate_*).

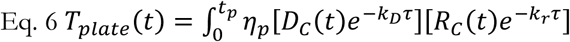

This framework produces several specific predictions. First, increasing antibiotic concentration accelerates the loss of donor or recipient conjugation competence, progressively reducing the residual conjugation window (**Fig. 3B**), consistent with our earlier observations (**Fig. 2B**). Second, the model predicts a characteristic biphasic trajectory in measured transconjugant abundance, with an initial decline followed by recovery (**Fig. 3C**). This behavior arises because the higher rate of RP4 transfers on solid surfaces(*22*, *28*, *32–34*) allows additional transconjugants to form during CFU plating, a contribution that progressively disappears as parent competence declines and growth of established transconjugants becomes dominant. Thus, the biphasic trajectory provides a distinct signature of declining conjugation competence during antibiotic exposure.

To directly test this prediction, we measured transconjugant dynamics during Tet and Rif exposure using the DA-wtRP4 donor and BWrifR recipient. Following conjugation, cells were exposed to constant 6x Rif and different concentrations of Tet (0, 5, 10, 15, 20 and 50x MIC), and sampled over time using CFU plating. As predicted, measured transconjugants exhibited a biphasic trajectory, with an initial decline followed by recovery; increasing Tet progressively extended and deepened this decline (**Fig. 3D**, points). To test whether these dynamics could be predicted from parent antibiotic responses alone, we independently fit donor and recipient growth rates and competence-loss parameters, as well as transconjugant growth rate (**Fig. S5**). Together, these parameters were sufficient to quantitatively predict the initial decline, subsequent recovery, and systematic shift in both the magnitude and duration of the decline with increasing [Tet] (**Fig. 3D**, lines). Thus, the observed transconjugant dynamics can be explained by the changing contribution of viable, conjugation-competent parent cells under antibiotic exposure.

Finally, we tested whether these competence-loss dynamics could explain the large antibiotic-specific differences in apparent background conjugation efficiencies. Using parameters independently estimated from background time courses for Kan and Carb (**Fig. S6**), together with the Tet and Rif parameters above, the model predicted the instantaneous residual conjugation potential over time for each antibiotic (**Fig. 3E**, *top*). Integration of these trajectories predicts substantially different cumulative opportunities for residual transfer, with Kan rapidly eliminating residual conjugation while Tet and Carb permit progressively longer transfer. Importantly, the model predictions closely reproduced the measured differences in conjugation efficiency across antibiotics (**Fig. 3E**, *bottom*). We note that for Carb, we assumed a 1-h delay in both antibiotic-mediated killing and thus loss of conjugation competence, consistent with the observed delay in its effect (**Fig. S6A**, *left*); relaxing this assumption did not meaningfully alter the predictions (**Fig. S6C**). Thus, drug-specific differences in the duration and magnitude of residual conjugation are sufficient to quantitatively explain the large antibiotic-specific differences in conjugation efficiency observed for the same multidrug resistance plasmid.

### Different strains exhibit unique residual conjugation windows

Our model predicts that the ability of residual conjugation to rescue a population from antibiotic selection depends strongly on population density. Intuitively, since conjugation requires encounters between donors and recipients, lower-density populations require a longer residual conjugation window to generate successful transconjugants, whereas higher-density populations can survive with a shorter window. Treating transfer as a stochastic process, we calculated the probability that at least one transconjugant lineage is established during the residual window, following a 1:1 donor recipient mixture ratio, assuming that a single successful transfer is sufficient for survival and that all dynamics occur in liquid culture. Using a Poisson model and defining survival at a 50% probability, a critical threshold emerges between populations likely or unlikely to survive (**Fig. 4A**, solid line). Importantly, the position and shape of this boundary are largely insensitive to the probability threshold (5% and 95% probabilities, dotted and dashed lines, respectively), demonstrating that the critical survival threshold shifts slightly, but the overall density-dependent relationship is preserved.

**Figure 4.**
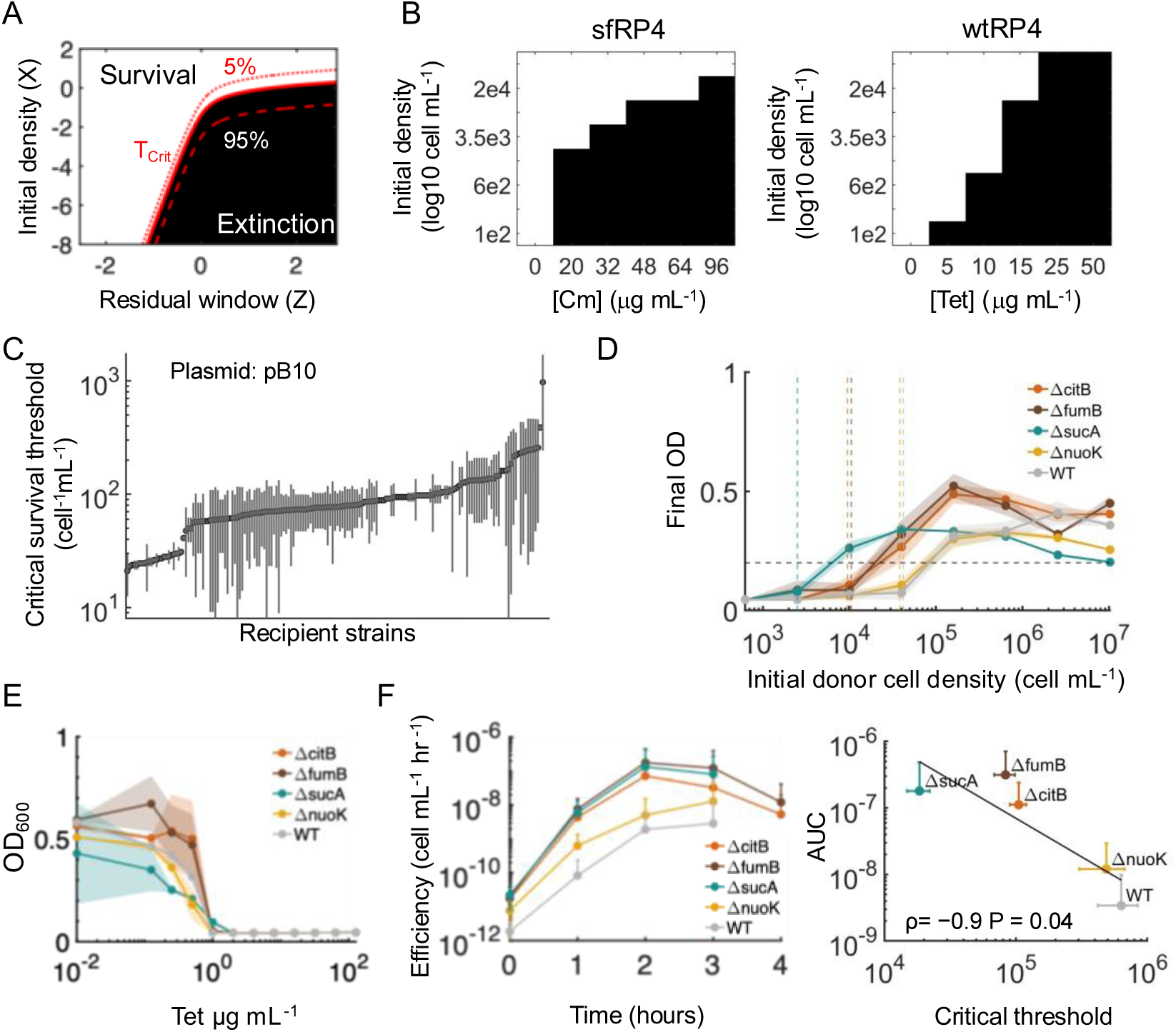
Different strains have unique residual conjugation windows. **A** Analytical prediction of the critical survival threshold as a function of initial transfer potential, (X=ηD_0_R_0_) (y-axis) varied by reducing density, and the rate of loss of transfer potential, (*Z* = −*μ_D_*_,*eff*_ − *μ_R_*_,*eff*_ + *k_D_*[*Rif*] + *k_R_*[*Tet*]) (x-axis), which incorporates antibiotic-dependent parent growth and loss of conjugation competence (see analytical derivation). Shading indicates conditions with <50% (black) or ≥50% (white) probability of generating at least one transconjugant; the solid, dashed, and dotted red lines indicate the 50%, 5%, and 95% critical survival probability threshold (T_Crit_). **B** Experimental validation of the critical survival boundary. Donor-recipient pairs MG-sfRP4/BWrifR (*left*) and DA-wtRP4/BWrifR (*right*) were mixed immediately before dilution into 96-well plates containing 6x Rif and increasing concentrations of Cm (0, 3, 5, 8, 10, 16x) or Tet (0, 5, 10, 15, 25, 50x), respectively. Following a 24 hour incubation at 37°C, OD₆₀₀ was measured. The highest replicate was excluded to account for spontaneous mutants in random wells, and the remaining replicates were averaged (n = 3). White and black boxes indicate survival and extinction, respectively. **C** Keio screen of 173 recipient strains using the pB10 donor. Parents were resuspended in M9CAG, mixed 1:1, incubated for 1 hour at 25°C. Cell mixtures were diluted 1:20 and subsequently serially diluted 1:4 in LB media containing 15µg/mL Tet and 25µg/mL Kan, to generate 6 initial densities being tested. After 24 hours at 37°C, the optical density at 600nm was measured. The lowest initial cell density that permitted growth was defined as the critical survival threshold. Growth was defined as OD > 0.2. Recipients are ordered from lowest to highest critical survival threshold and the complete list can be found in **Table S5**, attached separately. **D** Critical survival thresholds for selected Keio strains *(ΔcitB, ΔfumB, ΔsucA, ΔnuoK*) and BW WT (BW25113 carrying pLumi to provide Kan resistance). Strains were remeasured using the same assay but a higher resolution density gradient that started at ∼10^7^ cells per well, followed by 1:4 serial dilutions to ∼600 cells per well (8 densities total; n = 18). Experiments were conducted in triplicate with 6 biological replicates each; standard error of all 18 replicates is shown as shaded regions. The vertical dashed lines denote the critical donor density estimated from the mean growth curve, which is the average highest density that falls below the growth threshold of 0.2. The black dashed horizontal line denotes the OD threshold of 0.2. **E.** Tet MICs for selected Keio strains from (D) and BW WT. Strains were diluted 10,000-fold and then exposed to concentrations: 128, 64, 32, 16, 8, 4, 2, 1, 0.5, 0.25, and 0.125µg/mL, along with a no-drug control (12 total). The optical density was measured at 600nm after 24 hours of growth, and the MIC was determined based on the concentration that completely inhibited growth (OD < 0.1). The MIC was 1µg/mL for all strains. **F** Background time courses for Keio strains from (D) and BW WT (*left*). Time courses were performed using the standard protocol with parents each diluted 80-fold into LB. Backgrounds were calculated as efficiency at every time point, which was the transconjugant CFU divided by the donor times the recipient CFU. Area under the curve (AUC) was calculated from each background efficiency time course (*left*) and plotted against the corresponding critical survival threshold calculated directly from averaging the threshold-crossing densities across replicates (*right*). AUC was negatively correlated with critical survival threshold (Spearman, ρ = -0.9, P = 0.04).

We first experimentally tested the predicted dependence of the critical density on the residual conjugation window. We focused initially on the background condition, where no transconjugants are present before antibiotic exposure and survival therefore requires conjugation to occur entirely during antibiotic treatment, consistent with our model setup. Donors and recipients were mixed across a range of starting densities and exposed together in liquid culture to increasing Tet (for wtRP4) or Cm (for sfRP4) concentrations, which progressively reduce the residual conjugation window (e.g., **Fig. 2B**). As predicted, increasing either antibiotic shifted the critical survival threshold toward higher starting densities (**Fig. 4B**), confirming that populations with a more restricted opportunity for residual conjugation require more donor and recipient cells to survive. A similar density-dependent shift was also observed when transconjugants were already present before antibiotic exposure (e.g., mixture assay) (**Fig. S7**), highlighting that this relationship extends beyond conditions in which survival is strictly dependent on *de novo* transfer during treatment.

Together, these results suggest that the residual conjugation window can determine the population density required for plasmid-mediated survival under antibiotic selection. We therefore reasoned that biological factors beyond antibiotic concentration/identity that alter residual conjugation should similarly shift this critical survival threshold. To test this, we systematically perturbed the recipient genotype using the Keio(*35*) collection of *E. coli* single-gene knockout mutants. Since the Keio strains encode for Kan resistance, we used the plasmid pB10 belonging to the same incompatibility group as RP4 (incP) and was previously confirmed to exhibit residual conjugation (**Fig. S8**).

We screened 173 recipients from the first two Keio plates and defined the critical threshold as the lowest starting donor density supporting growth (OD > 0.2) (complete list of Keio gene knockouts in **Table S5** attached separately). Critical survival thresholds varied broadly across recipients, indicating substantial strain-to-strain variation in survival under these conditions (**Fig. 4C**). We selected four strains spanning the survival threshold range for further analysis: Δ*sucA*, Δ*nuoK*, Δ*citB* and Δ*fumB* (**Fig. 4D**). These genes affect distinct aspects of central metabolism and energy production, providing a functionally related set of mutants with distinct survival phenotypes. Specifically, while Δ*sucA*, which disrupts carbon oxidation and NADH production within the central TCA cycle, exhibited a low survival threshold, Δ*nuoK*, which impairs downstream respiratory electron transport, exhibited a high threshold. In contrast, Δ*citB* and Δ*fumB*, which affect more condition-dependent aspects of citrate and fumarate metabolism, respectively, exhibited intermediate thresholds. The WT control (BW25113 carrying pLumi for Kan selection) also exhibited a relatively high survival threshold.

According to our model, differences in the critical survival thresholds could either be explained by differences in antibiotic sensitivity, residual conjugation windows, or both. Indeed, while all strains exhibited the same MIC of 1 µg/mL to Tet (**Fig. 4E**), each recipient strain exhibited distinct residual conjugation dynamics (**Fig. 4F**, *left*), differing in both the magnitude and duration of continued transfer; we note that the residual window is expressed as conjugation efficiency instead of absolute transconjugant abundance to account for differences in donor and recipient population sizes over time.

Since the critical survival threshold should depend on the cumulative opportunity to generate transconjugants during antibiotic exposure, we summarized each residual conjugation window by calculating the area under its conjugation efficiency curve (AUC). This metric captures both how much conjugation remains possible and how long that capacity persists following antibiotic exposure. Strikingly, residual conjugation AUC was strongly negatively correlated with the independently measured critical survival threshold (Spearman, ρ = -0.9, P = 0.04; **Fig. 4F**, *right*). Recipients that sustained greater conjugation over the course of antibiotic exposure could support plasmid-mediated survival from lower starting densities, whereas recipients with smaller residual conjugation windows required higher initial densities. Thus, recipient-dependent differences in residual conjugation directly predict where the critical survival threshold emerges under antibiotic selection

## Discussion

Here, we show that antibiotics selecting for resistance genes encoded on the same multidrug-resistant plasmid can produce substantially different outcomes, with observed conjugation efficiency varying by nearly five orders of magnitude across experimental conditions. Much of this variation arises from an antibiotic-specific residual conjugation window, during which donors and recipients remain capable of generating transconjugants for several hours after bacterial growth has been inhibited. These findings demonstrate that growth inhibition does not necessarily prevent plasmid transfer and that the antibiotic used for selection can substantially influence the continued dissemination of multidrug resistance.

A simple quantitative model explains these differences through antibiotic-dependent loss of donor and recipient conjugation competence, recapitulating the observed conjugation efficiencies and predicting transconjugant dynamics over time. Under bactericidal conditions, greater parent killing was associated with reduced residual conjugation. However, under bacteriostatic conditions, conjugation competence progressively declined even after net population growth had stopped, revealing dynamics that cannot be explained by growth alone. Consistent with this observation, another study demonstrated that recipients could acquire tetracycline resistance through F-plasmid transfer during tetracycline exposure(*36*). Together, these results suggest that growth inhibition and loss of conjugation capacity do not necessarily occur simultaneously, and the residual window between these processes determines the opportunity for continued plasmid transfer under antibiotic selection.

The persistence of conjugation after growth inhibition suggests that the physiological requirements for plasmid transfer differ from those required for bacterial proliferation. Although conjugation depends on energy availability, DNA processing, and functional transfer machinery, these processes may remain active even when cells can no longer grow or divide. The distinct mechanisms of antibiotic action may therefore determine how long these functions remain intact following exposure. For example, bactericidal antibiotics such as carbenicillin and kanamycin can induce energy-consuming metabolic responses associated with cellular damage and stress(*37–39*), whereas bacteriostatic antibiotics such as tetracycline may preserve metabolic resources needed to sustain conjugation. However, the distinct residual conjugation windows observed for carbenicillin and kanamycin suggest that the timing and extent of these metabolic disruptions likely contribute to how long cells remain capable of plasmid transfer, beyond the distinction between bactericidal or bacteriostatic classification.

Our findings provide a mechanistic explanation for a long-recognized feature of conjugation assays, namely that measurements can reflect ongoing mating on selective agar in addition to during the intended mating period, particularly at high densities(*28*, *31*, *40*, *41*). Our results extend this observation by demonstrating that the extent of continued transfer also depends strongly on the selection antibiotic, such that measured transconjugant abundance can include antibiotic-specific transfer occurring after plating. Consistent with previous studies showing that cell density influences conjugation during selecting plating(*28*, *40–42*), our model predicts that lower-density populations require longer residual conjugation windows to generate successful transconjugants. Experimentally, increasing antibiotic concentration increased the minimum population density required for survival. Thus, beyond measured transfer efficiency, differences in the residual conjugation window may also influence the likelihood that plasmid-mediated resistance becomes established within a population.

We found that the relationship between population density and survival also depends strongly on recipient genotype. Indeed, critical survival thresholds varied substantially across 173 recipient strains and correlated strongly with residual conjugation measured in selected recipients, despite similar antibiotic susceptibility. Interestingly, the four recipients selected for further characterization all carried metabolic mutations but exhibited large differences in survival thresholds. This raises the possibility that continued conjugation depends on specific metabolic pathways or energetic requirements that remain functional following growth inhibition, although the underlying mechanisms remain to be determined. Extending our initial screen across the remaining Keio mutants could reveal pathway-level determinants of residual conjugation and help distinguish the metabolic requirements for continued plasmid transfer from the antibiotic-specific effects on these processes.

These dynamics have important implications for the co-selection of multidrug-resistant plasmids. Resistance genes encoded on the same plasmid are genetically linked, and selection for any one determinant can maintain the entire resistance element. However, epidemiological and experimental studies have repeatedly observed asymmetric co-selection, where different antibiotics selecting for the same linked resistance determinants produce different associations or persistence outcomes(*43*, *44*). Our results identify residual conjugation as one mechanism that could contribute to this asymmetry; antibiotics that select for the same plasmid can differ in how long they permit continued transfer into new hosts. Thus, co-selection may depend not only on the selective advantage of existing plasmid carriers, but also on the opportunity for continued horizontal transfer. Identifying the cellular mechanisms that govern residual conjugation could help explain antibiotic-specific patterns of co-selection and reveal new targets for developing antimicrobial strategies that limit ongoing plasmid transfer and the spread of resistance.

## Materials and Methods

### Strains, media, and growth conditions

In all cases, experiments were initiated from single colonies picked from antibiotic-containing agar plates (BD Difco, cat #DF0445076), inoculated in 2mL Luria-Bertani (LB) media (BD Difco, cat#DF0446075), and incubated overnight at 37°C for 16 hours shaking at 250rpm. Where applicable, LB was supplemented with specific antibiotics for selection: 15µg/mL for tetracycline (cat#B21408), 50µg/mL for rifampicin (cat#R0079), 25µg/mL for kanamycin (cat#61129-0050), 30µg/mL for chloramphenicol (cat#C0378-25G), 50µg/mL for gentamicin (cat#G1914-5G) and 100µg/mL for carbenicillin (cat#C1389-5G). In all cases, a colony-free LB negative control was included, and an experiment was discarded if growth was observed. A list of all strains and plasmids used in this study, along with all corresponding antibiotics used for selection can be found in **Table S2** and **Table S6.** Unless otherwise noted, all agar plates were incubated at 30°C until the end of day and then moved to 37°C for 16 hours, and all experiments were conducted using a minimum of three biological replicates.

### Minimum Inhibitory Concentration (MIC)

Single colonies of each strain were incubated for 16 hours overnight at 37°C, shaking at 250 rpm, supplemented with corresponding antibiotics when appropriate, as described above. Cells were diluted 10,000 times and added to a 96-well titer plate. Tetracycline, carbenicillin and kanamycin were added at the following concentrations: 128, 64, 32, 16, 8, 4, 2, 1, 0.5, 0.25, and 0.125 µg/mL, along with a no-drug control (12 total). Plates were then sealed with both a paper (AeraSeal, cat#BS-25) and a plastic membrane (Breathe-Easy, cat#Z380059-1PAK), and then incubated at 37°C shaking at 250 rpm for 24 hours. The optical density was measured at 600nm on a Tecan infinite Mplex plate reader. MIC was defined as the lowest antibiotic concentration that completely inhibited detectable growth (OD < 0.1). A lower OD threshold here is chosen to increase the sensitivity to growth to determine the most appropriate antibiotic inhibitory concentration. Specific MIC values for all strains are listed in **Table S1**, and MIC curves can be seen in **Fig. S2**.

### Conjugation assay

Individual donor and recipient clones were grown overnight, as described above. After 16 hours, cultures were spun down at 10,000rpm for 1 minute, resuspended 1:1 in M9CAG media, (M9CA medium broth powder, Bioworld cat#30621016, supplemented with 2mM MgSO_4_, 0.1mM CaCl_2_, and 0.4% w/v glucose), and incubated at room temperature for 5 minutes. For the *mixture condition*, equal volumes of donor and recipient were then combined, and donor-recipient mixtures were incubated for 1 hour in a 25°C stationary incubator to allow for conjugation. The conjugation mixture was then serially diluted in a 96-well plate, and 10μL from each dilution was spotted onto agar plates containing appropriate dual antibiotics or single parent selection antibiotics. For the *background condition*, resuspended parents were individually incubated for 1 hour in a 25°C stationary incubator in parallel; 5µL of recipient and donor serial dilutions were then each spotted onto the dual antibiotic plate in the same location (10µL total volume per spot). In both cases, donors and recipients were individually plated on their single antibiotic selective agar using the same protocol. See **Table S2** for the antibiotic used for each strain. Plates were incubated at 30°C until the end of day and then moved to 37°C for 16 hours of incubation. Dilutions that achieved single-colony resolution were used to record colony forming units (CFU); these were also used to calibrate plating densities for subsequent time course experiments. Conjugation efficiency of both the mixture and the background was calculated as the CFU of transconjugants formed divided by the CFU of donor and recipient cells along with the time of conjugation:

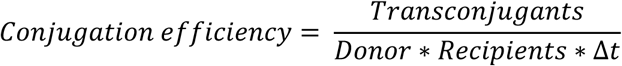

### Fluorescence and antibiotic resistance confirmation of transconjugants

A background mating experiment was conducted as described above using the BWrifR-wtRP4 donor and BWcmR recipient (**Fig. 1B**); a positive control (BWcmR recipient) and a negative control (BWrifR-wtRP4 donor) were also plated at the same time. Following 16 hours of growth on agar, 13 suspected transconjugant colonies from the dual antibiotic plates were picked and resuspended into 3 separate wells per colony (i.e., in triplicate) using LB containing 15µg/mL Tet and 31µg/mL Cm. 3 colonies of parent controls from single antibiotic plates were picked and resuspended in triplicate into LB containing their selective antibiotic of 15µg/mL Tet for donors and 31µg/mL Cm for recipients. 96-well titer plates were then covered with both membranes and incubated for 16 hours in the 37°C shaking incubator at 250rpm. To confirm the antibiotic resistance phenotype of presumed transconjugants, their growth in dual antibiotics was measured by reading absorbance at 600nm for all wells in a Tecan mPlex plate reader; GFP fluorescence was also read at excitation 475nm and emission 515nm to verify the fluorescence identity of presumed transconjugants. To account for differences in growth, fluorescence was normalized to the optical density to obtain an average fluorescence per OD measurement per sample (**Fig. S3C**).

### Flow cytometry of MG-sfRP4

BWrifR, MG-sfRP4, and resulting transconjugants (e.g., BWrifR-sfRP4) were inoculated from a single colony and grown for 16 hours at 37°C with shaking at 250 rpm. Cells were resuspended 1:1 in M9CAG, and donors and recipients were mixed in a 1:1 ratio. All cells were incubated for 1 hour at 25°C in static conditions to allow cells to conjugate. Following incubation, cells were diluted 1:100 in 1x PBS and analyzed on the Sony SH800 flow cytometer cell sorter. Voltages were adjusted according to the controls. Data was analyzed using FloJo software.

### Background and transconjugant time courses

Single colonies of donor and recipient strains were grown overnight as described above. Each population was centrifuged for 1 minute at 10,000rpm and resuspended 1:1 in M9CAG media. For the background time courses (e.g., **Fig. 2**), individual parents were incubated for 1 hour at 25°C; for the transconjugant time courses (e.g., **Fig. 3**), equal volume of donor and recipient were mixed and incubated under the same conditions. Following the incubation, cells were diluted ∼150x for individual parents, or ∼75x for the mixture into 6mLs of LB media, along with the indicated antibiotic and fold MIC used noted in the text and figure legends (see **Table S1** for the MIC). Tubes were placed in a 37°C incubator shaking at 250rpm (time 0). At 0, 1, 2, 3, 4, and 6 hours, an 80µL aliquot was removed, serially diluted in sterile water, and 10µL was used for CFU spot plating (mixture), or 5µL each of donor and recipient as described above. LB agar plates containing appropriate antibiotics were warmed in a 30°C incubator for 1 hour prior to spot plating. As needed for increasing resolution, at 2, 3, 4, and 6 hours spread plating was also performed by adding 50µL of recipient and 50µL of donor on top of one another and then spread using ∼5 sterile glass beads. Plates were flipped upside-down and incubated at 30°C until the end of day, at which point they were moved to 37°C for 16 hours for CFU enumeration.

### Shifted residual conjugation window matrix and critical cell density calculation

All strains were grown as previously described and resuspended in M9CAG media. The donors, DA-wtRP4 and MG-sfRP4, and the BWrifR recipient were incubated at 25°C in static conditions for 1 hour according to the background or mixture conditions described above. Following incubation, the background condition donors and recipients were mixed 1:1 and immediately diluted along with the mixture condition ∼75x for the wtRP4 or ∼40x for sfRP4; different dilutions were used based on differences in observed conjugation efficiencies and therefore expected residual conjugation window ranges. When parents alone were tested (**Fig. S7**), half the volume was added to match the parent density present in mixtures. Cells were added to LB media containing 50µg/mL (6x) Rif and then serially diluted 2.5-fold for a total of 8 initial density conditions. Tet was added to the 96-well plate at concentrations of 0, 5, 10, 15, 25, and 50µg/mL (corresponding to an MIC fold of 0x, 5x, 10x, 15x, 25x, and 50x); Cm was added at 0, 20, 32, 48, 64, and 96µg/mL (corresponding to 0x, 3x, 5x, 8x, 10x, and 16x). Plates were covered with a plastic and paper membrane, and incubated at 37°C with shaking at 250 rpm for 24 hours. After 24 hours, the optical density was measured at 600nm. Experiments were performed in duplicate each with two separate clones, resulting in four total replicates. Across all replicates, the well with the maximum OD was excluded from all conditions for subsequent analysis to account for any spontaneous mutants that appeared. Data was then averaged across experiments and converted to a binary growth/no growth outcome based on the OD threshold of 0.2 (or 0.15 with a background of 0.5 subtracted). An OD threshold of 0.2 was used to identify substantial outgrowth after 24 hours of incubation while minimizing the classification of minimal growth or minor OD fluctuations as growth.

### Keio recipient strain assay

Plates 1 and 2 from the Keio collection were stored in duplicate in 25% glycerol. Using a 96-well pinner, all strains (e.g, recipients) were grown from glycerol into a 96-well plate containing 200μL LB with 25µg/mL kanamycin. Plates were incubated at 37°C with shaking at 250 rpm for 16 hours. The pB10 donor was incubated in a flask containing 30mL of LB supplemented with 15x Tet. After incubation, 200μL of the donor was transferred into every well of a 96-well plate. Both donor and recipient plates were centrifuged at 4,680rpm for 10 minutes, after which the supernatant was removed, and cells were resuspended in 200µL of M9CAG media. 50µL of donors and 50µL of recipients were mixed together and incubated at 25°C for 1 hour to allow cells to conjugate. Following conjugation, mixtures were diluted 1:20, followed by dilutions of 1:4 (50µL into 150µL) to achieve 6 total densities. Dilutions were done using a 96-well multichannel pipettor into LB Broth containing 25µg/mL Kan and 15µg/mL Tet. All plates were covered with plastic and paper membranes, and incubated for 24 hours at 37°C with shaking at 250rpm. The absorbance at 600nm was measured using the Tecan infinite mplex plate reader. Each experiment was conducted in duplicate on the same day and on at least 2 different days. To calculate the critical survival threshold, MATLAB was used to interpolate (interp1) where the critical threshold is between the cell density in which the OD is > or equal to a threshold of 0.2 and the next lowest cell density that falls below 0.2. For the Keio screening experiments, an OD cutoff of 0.2 was selected to prioritize the detection of robust outgrowth after 24 hours, rather than weak growth signals or minor fluctuations in optical density. For subsequent strain validation, individual strains (*ΔcitB, ΔfumB, ΔsucA, ΔnuoK*, BW WT and the donor pB10 were streaked onto agar plates; all knockouts were verified via PCR (primers listed in **Table S8**). Critical survival thresholds were re-measured for all 5 strains following the same protocol with the following modifications: densities were initiated at an initial donor CFU of ∼10,250,000 cells per well followed by 1:4 dilutions until ∼600 cells per well (8 concentrations total); 6 replicates were included per strain and the experiment was repeated on three separate days. After setting up the experiment, all strains were diluted 1:10 in sterile water and spotted 10µl onto selective antibiotic agar plates for CFU enumeration. After plates were incubated at 37°C for 16 hours, the colonies were counted. Background time courses for the same five strains were conducted according to the standard protocol described above, with the following modifications: parents were diluted 80x into LB media to account for differences in observed conjugation efficiency of BW WT and pB10 donor. Background efficiencies over time were calculated as the transconjugant CFU divided by donor CFU times recipient CFU at that time point. To calculate the correlation between critical survival threshold and residual conjugation window, the area under the conjugation efficiency background curve was calculated from the background time course curves using the MATLAB function trapz and plotted against the average critical threshold. Spearman correlation was used to determine statistical significance.

### PCR verification of Keio strains

Candidate Keio knockout colonies were picked and resuspended in 20 µL sterile water, and 2 µL of the suspension was used as the DNA template for PCR with 2× Taq polymerase master mix according to the manufacturer’s instructions. BW WT was included as a control for each knockout strain. Knockout strains were verified by PCR using the junction-PCR strategy described for the Keio collection, in which primers within the kanamycin resistance cassette are paired with primers targeting the chromosomal regions flanking the disrupted gene(*35*). Specifically, the upstream forward primer (Up-Fwd) was paired with K1-Rev, and the downstream reverse primer (Down-Rev) was paired with K2-Fwd to verify the upstream and downstream cassette–chromosome junctions, respectively. PCR products were analyzed by 1% agarose gel electrophoresis. The observed amplicon sizes were compared with the expected sizes for each knockout. Primer sequences are provided in **Table S8**.

### Modeling residual conjugation dynamics

Donor and recipient densities were modeled using a concentration-dependent effective growth rate, *μ*_eff_ = *μ*/(1 + *KA*) − *aA*, where *μ* is the drug-free growth rate, *K*describes growth inhibition, *a* describes antibiotic-induced killing, and *A* is antibiotic concentration expressed as fold MIC (Eqs. 1-2). Drug-free growth rates for all populations (donors, recipients, and transconjugants), along with antibiotic-response parameters were fit sequentially from parent CFU time courses (**Fig. S5A-C, Fig. S6A**). The fractions of donors and recipients remaining conjugation-competent were modeled as exponential decays *F_D_* = *e*^−*A*^*^DkDt^*and *F_R_* = *e*^−*A*^*^RkRt^* (Eqs. 3-4). Rif donor and Tet recipient competence-loss rates were jointly fit to background-conjugation time courses using the independently determined parent dynamics (**Fig. S5D**), and these parameters were subsequently held fixed while fitting the corresponding Kan and Carb competence-loss rates (**Fig. S6B**). Transfer was modeled using the competent parent populations (Eq. 5) along with exponential growth. The complete five-state model (*D*, *R*, *F_D_*, *F_R_*, *T*) was solved numerically in MATLAB using ode45. Residual transfer on selective agar was calculated by integrating transfer from the competent parent populations over the plate incubation period, and total transconjugants were calculated as the sum of tube and plate contributions. All fitted parameters were then held fixed to predict time-resolved transconjugant abundance and antibiotic-dependent conjugation efficiencies. Point predictions used the fitted parameter values without multipliers. To propagate uncertainty in the single-time-point predictions, 100 samples of each drug-specific competence-loss rate were drawn from its fitted log-scale normal distribution, the plate integral was recalculated for each draw, and the resulting distribution was summarized across observations. The final model was used to predict time-resolved transconjugant abundance and conjugation efficiency for the Tet/Rif, Kan/Rif, and Carb/Rif conditions.

We derived an analytical solution for cumulative transfer across initial parent densities and Tet/Rif concentrations (see Supplementary Text), which was used to calculate the probability of plasmid-mediated survival across initial densities and antibiotic concentrations. Specifically, we treated conjugation as a Poisson process. For each combination of starting donor and recipient densities and Tet/Rif concentrations, the fitted model was used to calculate the expected cumulative number of successful transfer events over 16 h by integrating the instantaneous transfer rate from the conjugation-competent donor and recipient populations. The probability of generating at least one transconjugant was then calculated as *P* = 1 − *e*^−*λ*^, where *λ* is the expected cumulative number of transfer events. We assumed that each transfer event successfully established a transconjugant lineage and that a single established lineage was sufficient for plasmid-mediated survival. The heatmap reports this probability across the simulated parameter space, with the critical survival threshold defined by (P=0.5); boundaries corresponding to 5% and 95% probabilities were calculated analogously to evaluate the sensitivity of the predicted threshold to this definition. Full equations, fitting procedures, and analytical derivations are provided in the Supplementary Text.

## Acknowledgements

This study was funded by the National Institutes of Health award #1R35GM150871-01 (AJL), and the Royal Society University Research Fellowship URF\R1\231740 (TD).

## Data availability

All data generated in this study are provided in the Supplemental Information and/or is available upon request. The modeling code is available on GitHub and will be archived in Dryad digital repository (XX) upon publication.

## Author contributions

HDC conceived the research, designed and performed experiments, interpreted the data, conducted data analysis, and wrote the manuscript. GQ conceived the research, designed and performed experiments, interpreted the data, and edited the manuscript. AL performed experiments and interpreted the data. TD designed experiments, interpreted data, and edited the manuscript. CL provided the sfRP4 system used in experiments and edited the manuscript. AJL conceived the research, performed data analysis, interpreted the data, and wrote the manuscript.

