## Supplemental Information for "Antibiotic inhibition uncouples growth from horizontal gene transfer"

**This file contains:**
**Supplementary Text**
**Supplementary Figures S1:S8**
**Supplementary Tables S1:S9**

**Supplementary text**

We derived an analytical representation of the Tet/Rif tube-transfer model, that separates changes in donor and recipient abundance from loss of their ability to participate in conjugation following antibiotic exposure, to visualize how initial parent density and antibiotic-dependent loss of transfer potential jointly determine detectable conjugation. The calculation uses the same fitted parameters as the final workflow: donor and recipient drug-response parameters (μR, μD, aR, aD, Kr, kD), transconjugant growth rate (μT), Rif donor and Tet recipient competence-loss rates (kD, kR)), and the constant conjugation efficiency $\eta_0$. Values were based on the use of Tet and Rif as representative examples.

### Parent population dynamics

Donor and recipient abundance were described using a concentration-dependent effective growth rate:

$$\frac{dN}{dt} = \mu_{eff}\, N, \text{ where}$$

$$\mu_{eff} = \mu/(1 + KA) - aA$$

Here, N is parent CFU, $\mu$ is the drug-free growth rate, $A$ is antibiotic concentration expressed as fold MIC, $K$ describes concentration-dependent growth inhibition, and $a$ describes concentration-dependent killing. Thus, for immediate drug action,

$$N(t) = N_0 exp\left[\left(\frac{\mu}{1 + KA} - aA\right) t\right]$$

That is, for each donor and recipient, and corresponding antibiotic $A_R$ and $A_D$ the parent is sensitive to, were governed by the following equations:

Eq. S1 $\frac{dR}{dt} = \frac{\mu_{R\_eff} R}{A_D + K_R} - A_D a_R R$

Eq. S2 $\frac{dD}{dt} = \frac{\mu_{D\_eff} D}{A_R + K_D} - A_R a_D D$

Drug-free D and R growth rates were estimated by linear regression of log-transformed CFU during exponential growth (**Fig. S7A**). Drug-response parameters were then fit to the corresponding parent time courses by minimizing residuals between observed and predicted CFU. Rifampicin (Rif) and tetracycline (Tet) responses were represented primarily by $K$ (**Fig. S7B**), whereas kanamycin (Kan) and carbenicillin (Carb) responses used the fitted killing term $a$. Finally, we assumed that Carb killing only occurred after a 1-h delay, consistent with our data (**Fig. S8A**).

### Loss of conjugation competence

We independently modeled the fraction of each parent population that remained competent for conjugation. For Rif-exposed donors ($A_R$)) and antibiotic-exposed recipients (Tet, Carb, or Kan, $A_D$),

$$\text{Eq. S3: } \frac{dF_D}{dt} = -A_R\, k_D\, F_D$$

$$\text{Eq. S4: } \frac{dF_R}{dt} = -A_D\, k_R\, F_R$$

giving

$$F_D(t) = exp(-A_R\, k_D\, t), \quad F_R(t) = exp(-A_D\, k_R\, t)$$

The effective conjugating populations are therefore $D_C = DF_D$ and $R_C = RF_R$. The Rif donor and Tet recipient competence-loss constants, $k_R$ and $k_D$ respectively, were jointly fit to background-conjugation time courses using the independently fitted parent trajectories (**Fig. S7D**). These parameters were then held fixed while fitting the corresponding recipient competence-loss constants for Kan and Carb. All competence-loss constants were optimized in log space to constrain them to positive values.

### Transconjugant formation

Transfer in liquid culture was modeled using traditional bimolecular kinetics, proportional to the conjugation-competent donor and recipient populations, with newly formed transconjugants growing at the independently measured adapted growth rate $\mu_T$:

$$\text{Eq. S5: } \frac{dT_{tube}}{dt} = \mu_T\, T + \eta\, D\, R\, F_D\, F_R$$

where $\eta$ is the intrinsic conjugation efficiency kinetic rate constant. Eqs. S1–S5 were solved numerically to simulate time-resolved transconjugant abundance during antibiotic exposure in liquid culture. At each experimental sampling time, additional residual conjugation occurring after plating was calculated from the same competent parent populations and fitted competence-loss rates by integrating the transfer reaction over the plate incubation period:

$$\text{Eq. S6: } T_{plate}(t) = \int_0^{t_p} \eta_p [D_C(t)e^{-k_D\tau}][R_C(t)e^{-k_r\tau}]$$

where $\eta_p$, is the intrinsic conjugation efficiency kinetic rate constant on an agar plate. ($t_p$) was set to 16 h, consistent with our experimental growth procedure. We assumed that antibiotic-dependent rates of parent viability and competence loss remained unchanged after plating, while the conjugation coefficient was increased 100-fold to account for the enhanced transfer of IncP plasmids on solid surfaces reported previously(*22*, *28*, *31*, *41*). The total predicted transconjugant population at each sampling time was therefore calculated as the sum of transconjugants generated during liquid exposure and following plating ($T=T_{plate}+T_{tube}$). The fitted parent growth and competence-loss parameters were subsequently held fixed to predict time-resolved transconjugant abundance and antibiotic-dependent apparent conjugation efficiencies without fitting these measurements directly.

### Analytical form and interpretation

To obtain an interpretable analytical form, we considered the inhibitory regime used for the density-threshold analysis, in which donor and recipient growth were set to zero and their abundance declined through the antibiotic-dependent killing terms. For Tet and Rif,

$$D(t) = D_0\, exp(-a_R\, Rif\, t)$$
$$R(t) = R_0\, exp(-a_D\, Tet\, t)$$

Combining these expressions with the competence fractions gives

$$\eta D R F_D F_R = X\, exp(-Zt)$$

expressed as the instantaneous tube-transfer rate:

$$I(t) = \eta D(t) F_D(t) R(t) F_R(t) = X e^{-Zt}$$

where

$$X = \eta\, D_0\, R_0\, F_{D0}\, F_{R0}$$
$$Z = Rif(a_D + k_D) + Tet(a_R + k_R)$$

X therefore captures the initial opportunity for conjugation, determined by starting parent densities and competence, whereas Z captures the combined antibiotic-dependent loss of that opportunity through parent killing and loss of competence. Both competence fractions are initialized to one; therefore $X = \eta D0R0$, has units of CFU $h^{-1}$ and is the initial rate of successful transfer before parent dynamics and competence loss reduce that rate. Z has units of $h^{-1}$ and summarizes the combined loss of transfer potential. The transconjugant equation reduces to

$$dT/dt = \mu\_T\, T + X\, exp(-Zt)$$

For the deterministic heatmap, transconjugants were initialized at zero and allowed to grow at the independently fitted adapted-transconjugant rate $\mu_T$, evaluated at 15-fold Tet. Integrating the differential equation for Transconjugants yields:

$$T(t) = \frac{X}{Z + \mu_T} [exp(\mu_T t) - exp(-Zt)]$$

This expression shows directly how the final transconjugant population reflects competition between the initial opportunity for transfer, X, antibiotic-driven loss of conjugation potential, Z, and subsequent transconjugant growth $\mu_T$.

### Critical-density prediction

The analytical solution predicts the initial transfer opportunity required to reach a specified transconjugant threshold T_crit at time t. Rearranging gives

$$X_{crit} = T_{crit} \frac{Z + \mu_T}{[exp(\mu_T t) - exp(-Zt)]}$$

We tested this prediction by simulating the full ODE model across combinations of initial donor and recipient densities and Tet/Rif concentrations. The analytical X_crit boundary was then overlaid on the numerical phase diagram assuming an observed cutoff of 1e8 based on the OD limit of detection, providing a direct comparison between the simplified analytical prediction and the full model.

The stochastic companion heatmap treats established transfer events within a single well as an inhomogeneous Poisson process. A transfer event at time s contributes exp[μT(tend − s)] descendants at the endpoint. The latest transfer time that can meet a detection threshold N_detect is:

$$t_{detect} = t_{end} - \frac{log(N_{detect})}{\mu_T}$$

With P_establishment as the probability that a transfer event establishes, the expected number of established detectable events is:

$$\Lambda = p_{establishment} \int_0^{t_{detect}} Xe^{-Zt}\, dt = p_{establishment} X \frac{1 - e^{-Zt_{detect}}}{Z}$$

At Z = 0, the continuous limiting form Λ = P_establishment X t_detect was used. The probability that the individual well is positive is:

$$P(positive\ well \mid X, Z) = 1 - e^{-\Lambda}$$

For a chosen positive-well probability p, the corresponding boundary in X–Z space is:

$$X_{crit}(Z; p) = \frac{-log(1 - p)\, Z}{p_{establishment}(1 - e^{-Zt_{detect}})}$$

**Heatmap implementation.** Tet and Rif concentrations are each evaluated from 0 to 50-fold MIC in five-fold increments. Initial donor and recipient densities are each evaluated from $10^2$ to $10^7$ CFU, and their combinations are converted to X. The condition-specific Z values are resampled to a 500-point linear grid and X values to a 500-point logarithmic grid. The probability heatmap displays P(positive well | X, Z). The binary heatmap uses a 50% positive-well probability as its primary boundary, with 5% and 95% probability boundaries plotted as dashed curves.

**Default stochastic settings.** The current script uses tend = 16 h, P_establishment = 1, and N_detect = 1 descendant cell. Thus t_detect = 16 h and the positive-well probability represents the probability that at least one successful transfer occurs in that individual well by the endpoint. It does not pool or average transfer events across wells.

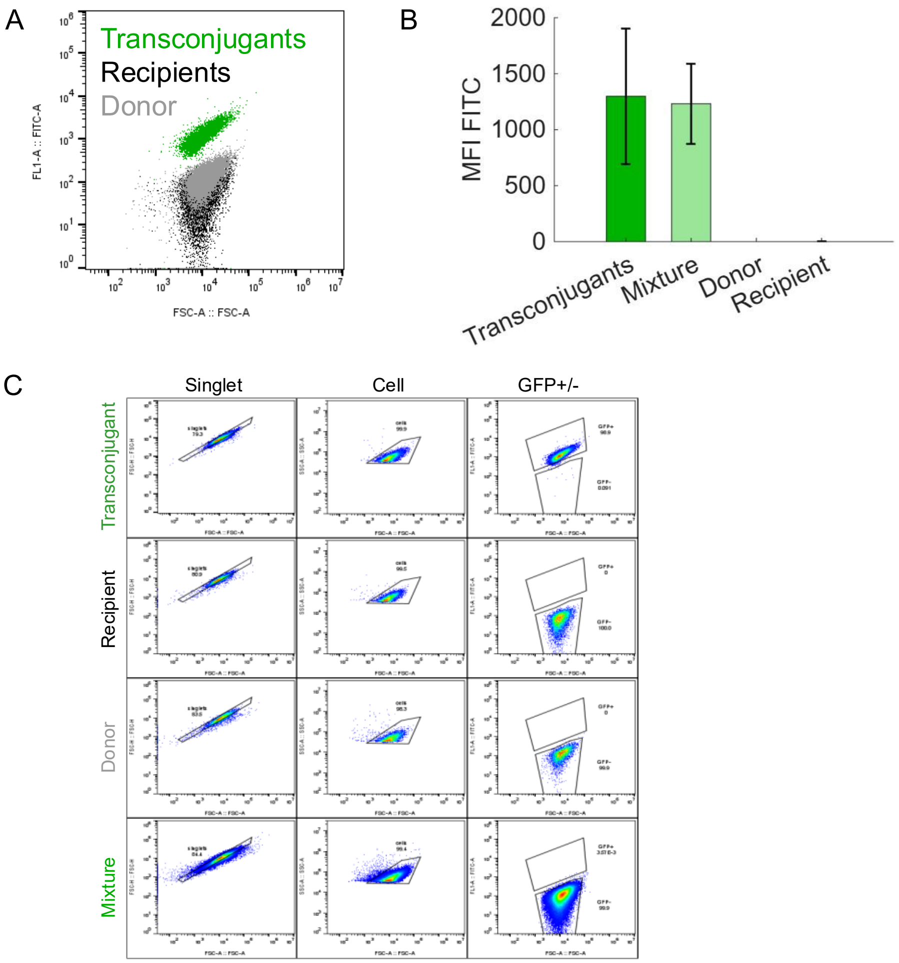


**Figure S1: Verification of sfRP4 transconjugants after the standard conjugation protocol**

A. Confirmation of sfGFP fluorescence: Pure populations of transconjugants (BWrifR-sfRP4), recipients (BWrifR), and donors (MG-sfRP4) were analyzed for FITC expression on the flow cytometer; all three populations are overlaid onto the same graph for visualization. Both the recipient (black population) and the donor do not have high FITC fluorescence expression but the transconjugants which encodes sfGFP is brightly expressed (green population).

B. Verification of transconjugants after conjugation: Donor MG-sfRP4 and recipient BWrifR were diluted in M9CAG, mixed together in a 1:1 ratio, and incubated statically for 1 hour at 25°C to allow for conjugation to occur. After conjugation, the donor, recipient, the conjugation mixture and a pure population of transconjugants were diluted 1:100 in 1x PBS and run on the flow cytometer. Both the transconjugants (green bar) and the mixture (light green bar) conditions show high FITC fluorescence The bar height represents the mean fluorescence intensity (MFI) of FITC and the error bars are the standard deviation.

C. Gating Strategy. Samples were first gated for singlets on FSC-Area and FSC-Height (*left column*), then for cells on FSC-Area and SSC-Area (*middle column*) and then the transconjugants and recipient cells were used to mark the GFP+ and GFP- gates cells (*right column*). Within the GFP+ gate, there were 30,398 cells of transconjugants, 9 cells of mixture transconjugants, 0 donor and 0 recipient cells.

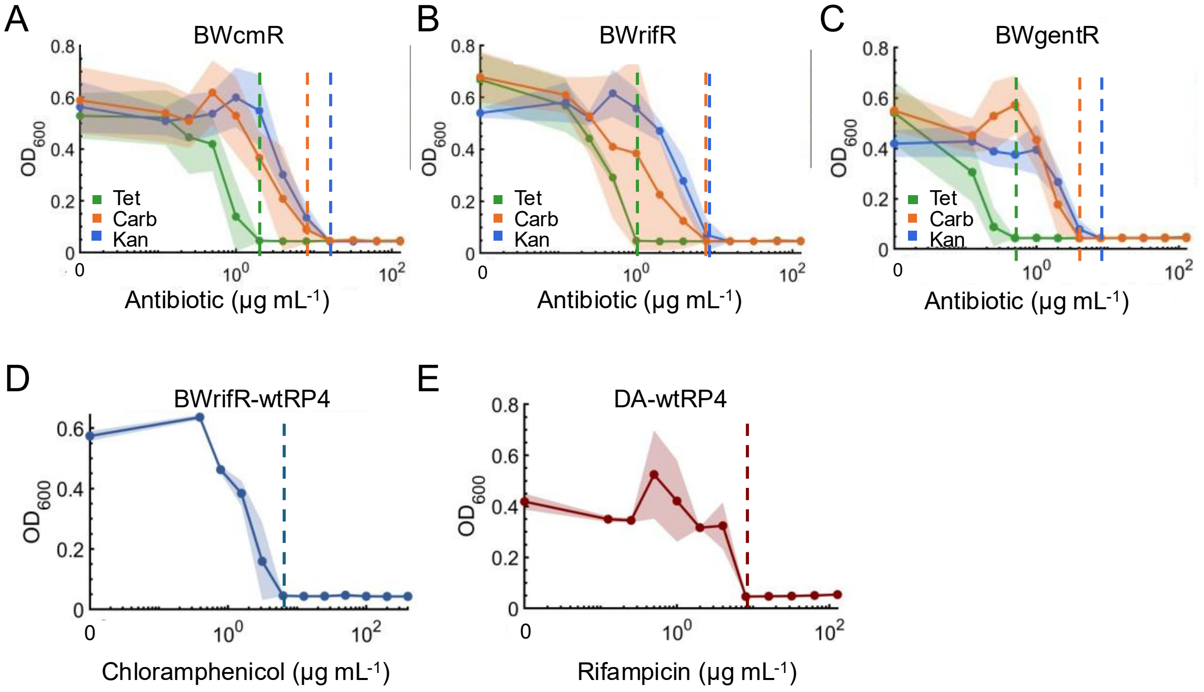


**Figure S2: Minimum Inhibitory Concentrations (MIC) of donors and recipients.**
In all cases, overnight cultures were diluted 10,000-fold in LB media, and experiments were conducted in biological triplicate. The MIC (dashed vertical line) is defined as the lowest concentration of antibiotic that fully inhibits growth, based on an OD < 0.1 Experiments were conducted in biological triplicate.

A. BWcmR Recipient: Tet (green), Kan (blue), or Carb (orange) was added at concentrations: 128, 64, 32, 16, 8, 4, 2, 1, 0.5, 0.25, 0,125, and 0μg/mL. The recipient strain BWcmR has an MIC for Tet at 2μg/mL, Kan at 16μg/mL and Carb at 8μg/mL. See **Table S1** for the full summary.
B. BWrifR Recipient: Tet, Kan, or Carb was added at concentrations: 128, 64, 32, 16, 8, 4, 2, 1, 0.5, 0.25, 0,125, and 0μg/mL. The recipient strain BWrifR has an MIC for tetracycline at 1μg/mL, kanamycin at 8μg/mL and carbenicillin at 8ug/mL.
C. BWgentR Recipient: Tet, Kan, or Carb was added at concentrations: 128, 64, 32, 16, 8, 4, 2, 1, 0.5, 0.25, 0,125, and 0μg/mL. The recipient strain BWgentR has an MIC for tetracycline at 0.5μg/mL, kanamycin at 4μg/mL and carbenicillin at 8ug/mL.
D. BWrifR-wtRP4 Donor: Chloramphenicol was added at: 400, 200, 100, 50, 25, 12.5, 6.25, 3.125, 1.56, 0.78, 0.39 and 0μg/mL. The MIC of BW25113rifR+RP4 to chloramphenicol is 6.25μg/mL.
E. DA-wtRP4 Donor: Rifampicin was added at: 128, 64, 32, 16, 8, 4, 2, 1, 0.5, 0.25, 0,125, and 0μg/mL. The MIC of DA-wtRP4 to rifampicin is 8μg/mL.

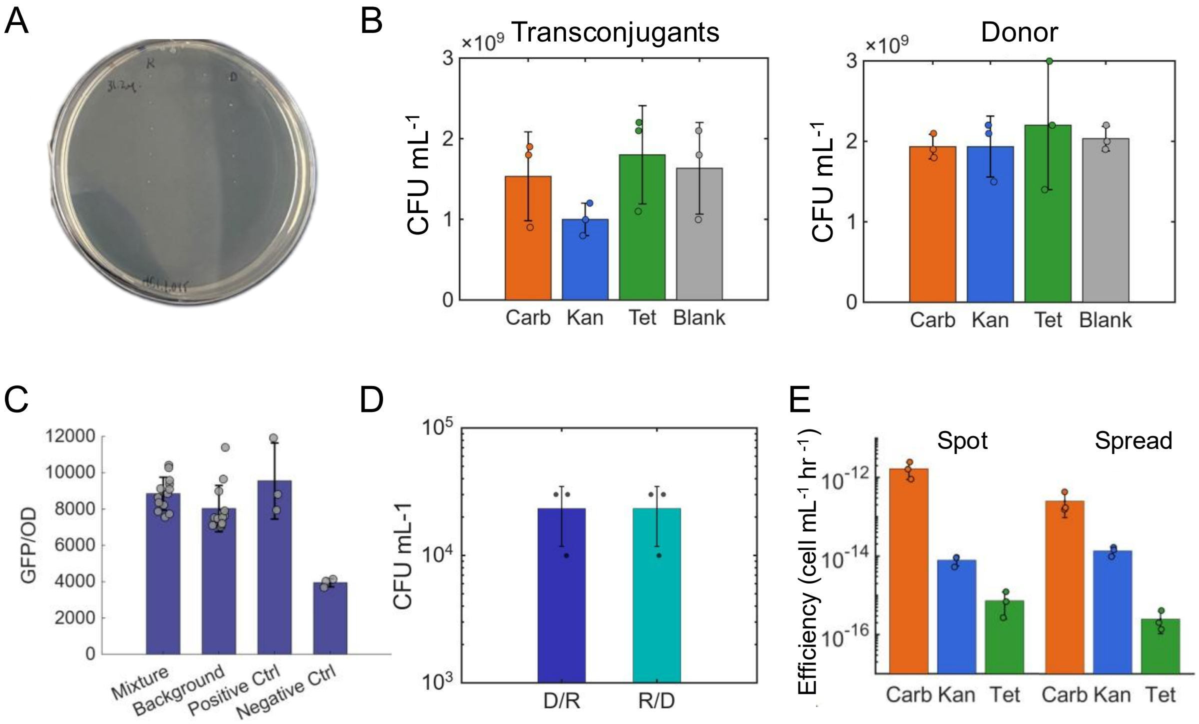


**Figure S3: Verification of transconjugant identity from the background condition (Fig. 1B). In all cases, experiments were conducted in biological triplicate.**

A. Parent controls. Donor BWrifR-wtRP4 and recipient BWcmR individually do not grow on the 15x Tet and 5x Cm dual drug plates.
B. Different antibiotics support equal growth of established transconjugants and donors. Overnight cultures of transconjugant strain BWcmR-wtRP4 (A) and donor strain BWrifR-wtRP4 (B) were diluted and plated onto three antibiotic-containing agar plates at 15x the MIC (30µg/mL Tet, 120µg/mL Carb, and 240µg/mL Kan). The blank plate was LB Agar without any antibiotics added. are both shown. In both cases, the growth on antibiotic plates was insignificantly different than growth on blank plates ($P > 0.9$, one-way ANOVA).
C. Transconjugant characterization. 13 colonies were picked from both the mixture and background conditions, grown in liquid LB containing dual selection antibiotics (5x Cm and 15x Tet). Three colonies for each parent were picked and grown in their single selection antibiotic to be used as controls. The positive control was recipient BWcmR, which encodes for GFP, and the negative control was the donor BWrifR-wtRP4. Cells were measured after 16 hours of growth at 37°C shaking. The OD was measured at 600nm and the GFP fluorescence was read at excitation/emission 475/515nm. The y-axis shows the GFP fluorescence per OD to normalize to growth.
D. Plating order of the parent background does not impact transconjugant background. Plating the recipient first then the donor on top (left bar) vs the donor first onto the plate then the recipient on top (right bar) results in insignificantly different transconjugant yields (paired two-tailed t-test $P = 1$).
E. Plating method does not alter antibiotic trends. Spread plating 100µL of the background (50µL of donor + 50µL of recipient) yields similar trends of efficiency across drugs (one-way ANOVA $P = 0.02$) as the 10µL spot method (5µL of donor +5µL of recipient) (ANOVA, $P = 0.006$). The efficiency does differ for Carb between the plating method (two-tailed ttest, $P = 0.04$), but not for Kan ($P = 0.07$) or Tet ($P = 0.2$) ($n = 3$).

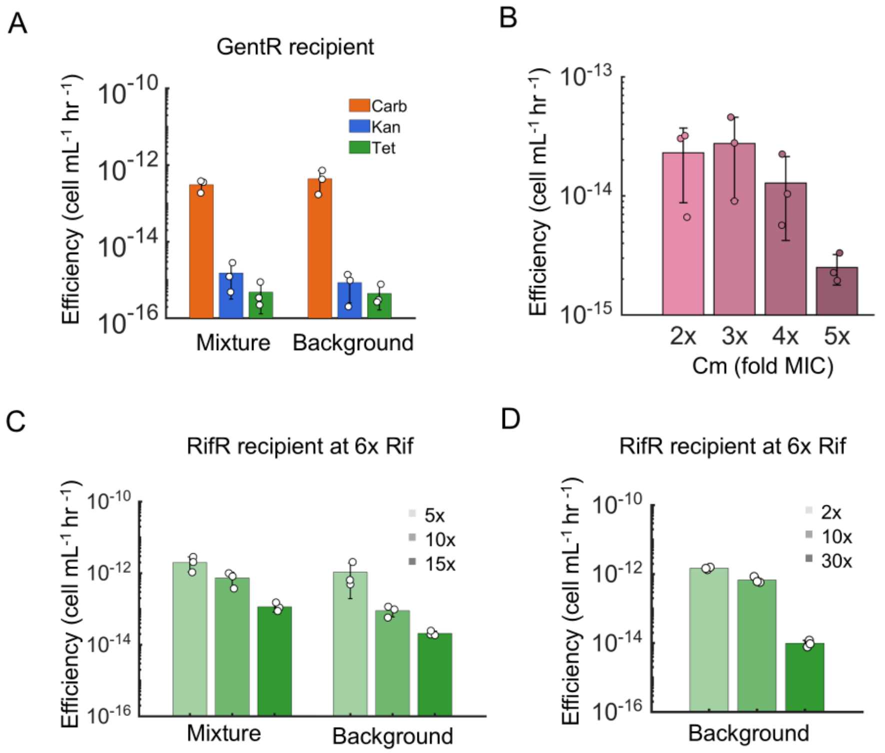


**Figure S4: Conjugation efficiency shows similar antibiotic and dose trends across recipient and antibiotic concentrations.**

A. The mixture vs background efficiencies of the GentR recipient under Carb, Kan, or Tet selection. The gentamicin-resistant recipient BWgentR and donor BWrifR-wtRP4, or their combination (e.g., mixture condition) were incubated statically at 25°C for 1 hour. Following incubation, 10µL of the conjugation mixture or 5 µL of each parent (background), was plated onto agar containing 15x the MIC of the plasmid-selecting antibiotic (7.5µg/mL Tet, 60µg/mL Carb, and 120µg/mL Kan) and 5x the MIC of gentamicin. Differences between the mixture and background efficiency were for all drugs is not significant (paired ttest with Bonferroni correction Carb: P = 1, Tet: P = 1, Kan P = 0.3 n = 3). However, responses differed significantly among the three selecting antibiotics for both the mixture (one-way ANOVA P = 5x10$^{-5}$) and the background (P = 7.5x10$^{-5}$) conditions.

B. Recipient antibiotic dose dependence. Recipient BWcmR and donor BWrifR-wtRP4 were conjugated under standard conditions for background and then spot plated onto agar plates containing either 2x, 3x, 4x or 5x the MIC of chloramphenicol (12.5, 18.7, 25, and 31µg/mL). With decreasing chloramphenicol doses, the transconjugant yield increases (Spearman ρ = -0.73, P = 0.007). Experiments were conducted in triplicate. Bar heights represent the average, and the error bars denote the standard deviation.

C. The mixture vs background efficiencies of the RifR recipient under higher Rif (6x) and doses of Tet selection. The Rif-resistant recipient BWrifR and donor DA-wtRP4, or their combination were incubated statically at 25°C for 1 hour. Following incubation, 10µL of the conjugation mixture or 5 µL of each parent (background), was plated onto agar containing 5x, 10x, or 15x the MIC of tetracycline (5, 10, 15µg/mL respectively) and 6x the MIC of rifampicin (50µg/mL). Mixture efficiency (*left*) decreased with increasing concentrations of Tet (Spearman ρ = -0.9, P = 0.001, n = 9), as did the background efficiency (*right*) (ρ = -0.9, P = 0.001, n = 9)

D. The background efficiencies of the RifR recipient under higher Rif (6x) at 2x, 10x, and 30x Tet selection. The rifampicin-resistant recipient BWrifR and donor DA-wtRP4, were incubated statically at 25°C for 1 hour. Following incubation, 10μL of the conjugation mixture or 5 μL of each parent (background), was plated onto agar containing 2x, 10x, or 30x the MIC of tetracycline (2, 10, 30μg/mL respectively) and 6x the MIC of rifampicin (50μg/mL). Pairs with **Fig.2B**, *right* to show dose dependence even at 6x rifampicin. Conjugation efficiency decreased with increasing concentrations of Tet (Spearman $\rho$ = -0.9, P = 0.001, n = 9).

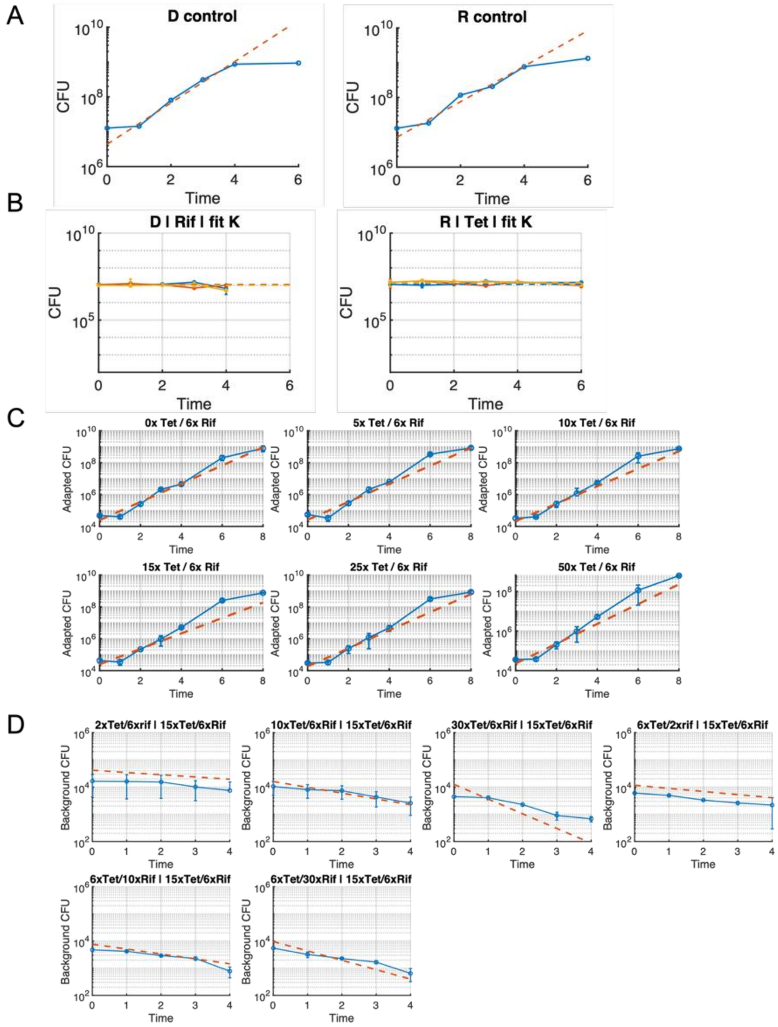


**Figure S5. Parameterization of parent growth and residual conjugation dynamics under Tet and Rif exposure.**

A. Drug-free donor and recipient growth controls to calculate baseline growth rates μ.
B. Donor and recipient CFU dynamics under Rif and Tet treatment, to estimate antibiotic-dependent *K* parameters. Colors indicate 5, 10, and 15x antibiotic concentration for blue, red, and yellow respectively.
C. Transconjugant growth dynamics across a range of Tet concentrations, consistent with **Fig. 3D**.
D. Background conjugation measured after progressively longer antibiotic exposure of the donor or recipient. Parent growth parameters estimated in panels **A-C** were held fixed, and the decline in the conjugation-competent donor and recipient fractions was fit to the background conjugation measurements to estimate the corresponding competence-loss parameters ($k_D$ and $k_R$). Blue line shows measured background transconjugant CFU and red dashed line shows model fits. Together, these independently parameterized parent growth and competence dynamics were subsequently used to predict transconjugant trajectories without fitting the transconjugant time-course measurements directly.

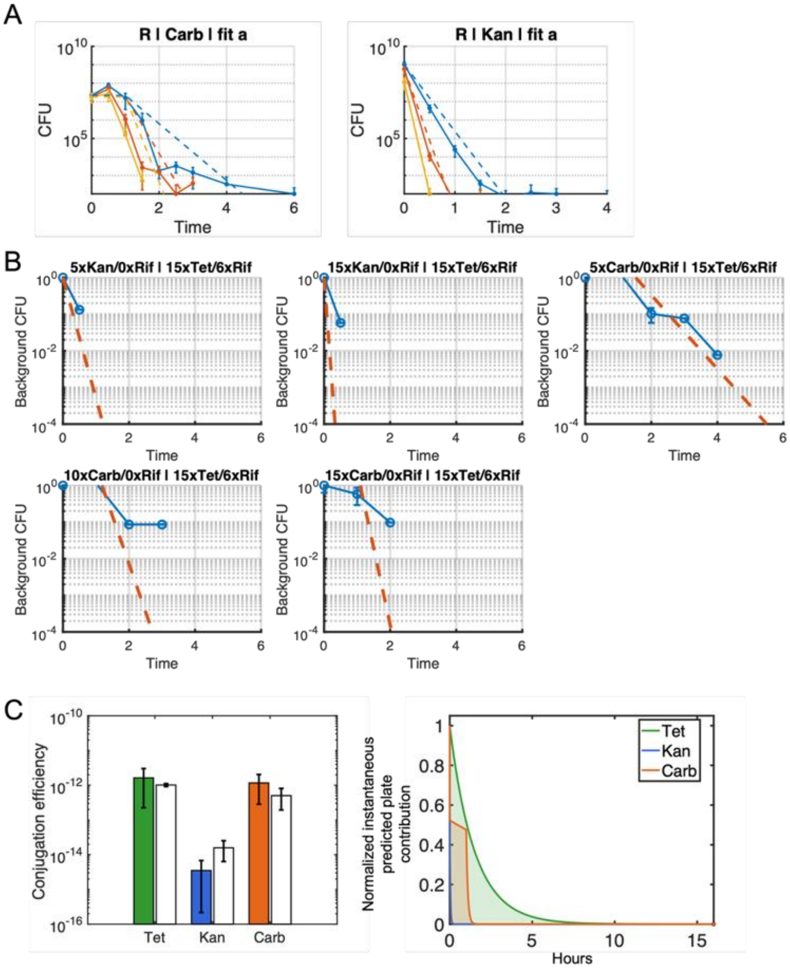


**Figure S6. Parameterization of parent growth and residual conjugation dynamics under Kan and Carb exposure.**

A. Recipients BWrifR were exposed to increasing concentrations of Carb (*left*) or Kan (*right*) of 5x, 10x, and 15x the MIC; CFU was measured hourly for up to 6 hours. Bactericidal killing rates ($a_{kan}$ and $a_{carb}$) were fit using all 3 concentrations together as described in the Supplementary text.

B. Background conjugation measured after progressively longer antibiotic exposure of the donor or recipient to varying Carb or Kan; the killing rate for $k_D$ (Rif) was assumed to be the same as from **Fig. S5**. All parameters were fit simultaneously.

C. The primary model assumed a one-hour delay in Carbenicillin killing, and thus a delay in conjugation inhibition as well. Relaxing this assumption by fitting the exponential killing for Carb and using immediate plate-based conjugation killing does not meaningfully change the results.

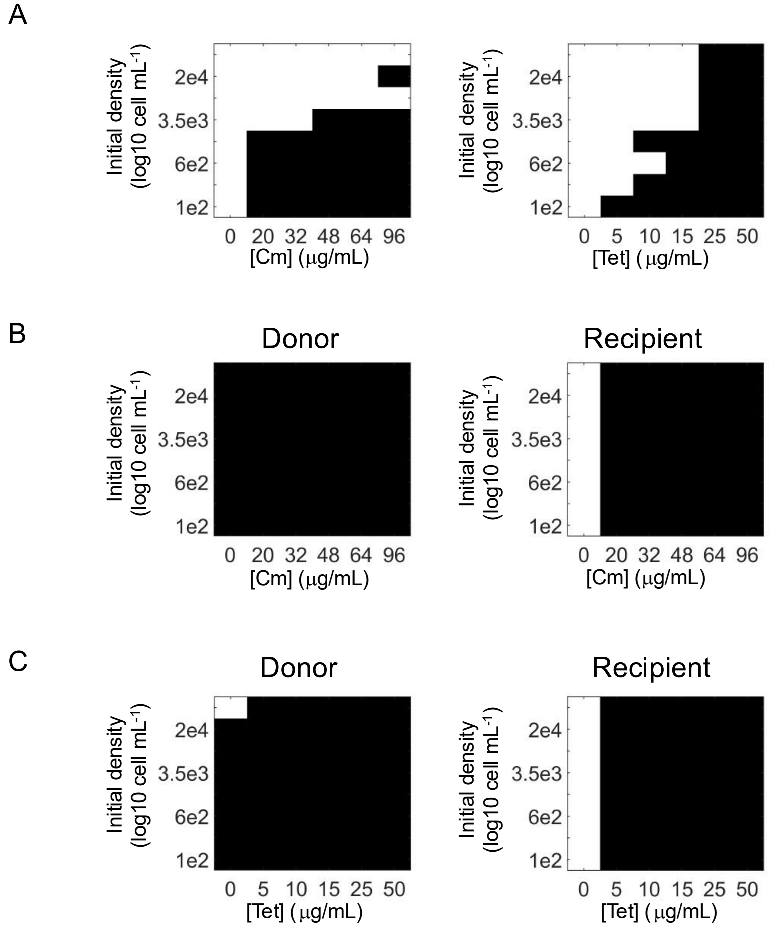


**Figure S7**: **Residual conjugation window experimental validation**. Residual conjugation window of two donors, under a range of selection antibiotic concentrations Cm or Tet, respectively. Tet was added to the 96-well plate at concentrations of 0, 5, 10, 15, 25, and 50 μg/mL, while Cm was added at 0, 20, 32, 48, 64, and 96 μg/mL. Cells were added to LB media containing 50ug/mL Rif and then serially diluted 2.5-fold for a total of 8 initial density conditions. Optical density was measured at 600nm after 24 hours of incubation. The highest value replicate was removed and then data was averaged across experiments and converted to a binary growth/no growth outcome based on the OD threshold of 0.2 (or with background subtracted the threshold is 1.5). Growth is indicated by white boxes, whereas no growth is indicated by black boxes. Experiments were performed in duplicate with two separate clones, resulting in four total replicates.

A. Mixtures of donors, MG-sfRP4 (A, *left*) and DA-wtRP4 (B, *right*), and recipients BWrifR were conjugated and then diluted in varying concentrations of their selection antibiotic, rifampicin and either chloramphenicol or tetracycline. Following incubation, the background condition donors and recipient were mixed 1:1 and immediately diluted along with the mixture condition ~75x for the wtRP4 or 42x for sfRP4; different dilutes were used based on differences in observed conjugation efficiencies, and therefore expected residual conjugation window ranges.
B. Parents MG-sfRP4 and BWrifR were incubated and diluted separately in chloramphenicol and rifampicin at consistent volumes to the Mixture (A) and background (**Fig.4B**) conditions. Half the volume was added to match the parent density present in mixtures
C. Parents DA-wtRP4 and BWrifR were incubated and diluted separately in chloramphenicol and rifampicin at consistent volumes to the Mixture (A) and background (**Fig.4B**) conditions. Half the volume was added to match the parent density present in mixtures

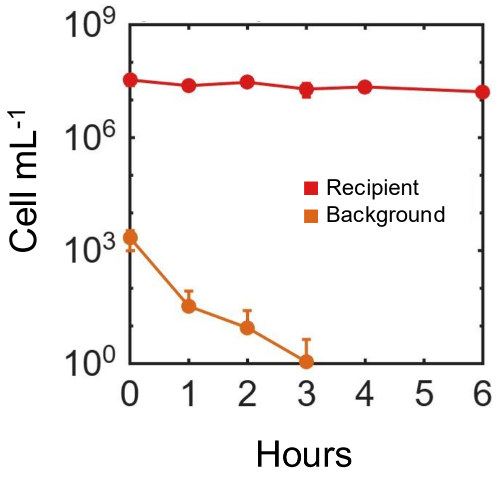


**Figure S8**: **BW WT with donor pB10 time course.**
BW WT and the donor pB10 were centrifuged for 1 minute at 10,000 rpm and resuspended 1:1 in M9CAG media. The individual parents were incubated for 1 hour at 25°C. The donor and recipient were diluted 80x into LB media and placed in a 37°C incubator shaking at 250 rpm (time 0). At 0, 1, 2, 3, 4, and 6 hours, an 80μL aliquot was removed, serially diluted in sterile water, and 10μL of recipient was used for CFU spot plating. Onto LB agar containing 15μg/mL tetracycline and 25μg/mL kanamycin, 5μL of recipients were plated then 5μL of donors were spotted right on top of the wet recipient droplet. At 2, 3, 4, hours 50μL of recipient and 50μL of donor were mixed and spread using ~5 sterile beads to increase the resolution. Plates were incubated at 30°C until the end of day, at which point they were moved to 37°C for 16 hours. The calculated CFU/mL for both the background transconjugants (orange) and the recipient (red) is plotted over time (n = 6).

**Supplementary Tables**

**Table S1:** Complete table of each strain and its absolute value MIC (ug mL$^{-1}$) for each drug. NA denotes that the MIC for that antibiotic couldn't be measured, since it has resistance. A "—" indicates that that antibiotic wasn't measured for that strain.

| **Strain** | **Additional antibiotic** | Tet | Carb | Kan | Cm | Rif | Gent |
|---|---|---|---|---|---|---|---|
| BWrifR | NA | 1 | 8 | 8 | 6 | NA | 25 |
| BWcmR | 5X Cm | 2 | 8 | 16 | NA | — | — |
| BWgentR | 5X Gent | 0.5 | 4 | 8 | — | — | NA |
| MG | NA | 2 | NA | NA | NA | 8 | 8 |
| DA-wtRP4 | NA | NA | NA | NA | NA | 8 | — |
| BWrifR-wtRP4 | NA | NA | NA | NA | 6.25 | NA | — |

**Table S2:** Complete table of strains and plasmids used in this study.

| **Name** | **Strain** | **Genotype** | **Plasmid** | **Resistances (used for selection*)** | **Source** |
|---|---|---|---|---|---|
| DA | DA838 | DA32838 Eco galK::cat-J23101-dTomato | wtRP4 | tetracycline*, carbenicillin, chloramphenicol, kanamycin | Lopatkin *et al*, 2016 (*29*) |
| MG | MG1655 | MG1655 (ilvA::placIQ1-tetR-FRT) + sRP4-ptetRA-sfGFP-frt-kn-frt-delta-tetRA-frt-Cm-Frt | sfRP4 | chloramphenicol*, streptamycin, kanamycin | Generous gift from Christian Lesterlin |
| BWrifR | BW25113 | E. coli BW25113 (F- , D(araD-araB)567, DlacZ4787(::rrnB-3), λ–, rph-1, D(rhaD-rhaB)568, hsdR514, rpoBM) | | rifampicin | Ahmad *et al,* 2023 (*45*) |
| BWcmR | BW25113 | E. coli BW25113 (F- , D(araD-araB)567, DlacZ4787(::rrnB-3), λ–, rph-1, D(rhaD-rhaB)568, hsdR514) | pZE31-GFP | chloramphenicol | Lab stock |
| BWgentR | BW25113-pCRIMSONgentR | E. coli BW25113 (F- , D(araD-araB)567, DlacZ4787(::rrnB-3), λ–, rph-1, D(rhaD-rhaB)568, hsdR514) | | gentamicin | Lab stock |

| | | | | | |
|---|---|---|---|---|---|
| BWpZE31 transconjugants | BW25113pZE31-GFP | E. coli BW25113 (F- , D(araD-araB)567, DlacZ4787(::rrnB-3), λ–, rph-1, D(rhaD-rhaB)568, hsdR514) | pZE31-GFP + RP4 | chloramphenicol, tetracycline* | Lab stock |
| pB10 donor | BW25113rifR | E. coli BW25113 (F- , D(araD-araB)567, DlacZ4787(::rrnB-3), λ–, rph-1, D(rhaD-rhaB)568, hsdR514, rpoBM) | pB10 | tetracycline, streptamycin | Ahmad *et al,* 2023 (*45*) |
| BWrifR-wtRP4 | BW25113rifR | E. coli BW25113 (F- , D(araD-araB)567, DlacZ4787(::rrnB-3), λ–, rph-1, D(rhaD-rhaB)568, hsdR514, rpoBM) | wtRP4 | tetracycline*, rifampicin | Ahmad *et al,* 2023 (*45*) |
| Δ*sucA* | BW25113 b0726: Keio Plate 2 Well E12 | rrnB DElacZ4787 HsdR 514 DE(araBAD)567 D E(rhaBAD)568 rph-1 ΔsucA | | kanamycin | Baba *et al*, 2006 (*35*) genotype info from: https://shigen.nig.ac.jp/ecoli/strain/resource/strainGeneMutant/list?strainNo=ME9062 |
| Δ*citB* | BW25113 b0620: Keio Plate 4 Well C9 | rrnB DElacZ4787 HsdR 514 DE(araBAD)567 D E(rhaBAD)568 rph-1 ΔcitB | | kanamycin | Baba *et al*, 2006 (*35*) |
| Δ*fumB* | BW25113 b4122: Keio Plate 4 Well H7 | rrnB DElacZ4787 HsdR 514 DE(araBAD)567 D E(rhaBAD)568 rph-1 ΔfumB | | kanamycin | Baba *et al*, 2006 (*35*) |
| Δ*nuoK* | BW25113 b2279: Keio Plate 4 Well B4 | rrnB DElacZ4787 HsdR 514 DE(araBAD)567 D E(rhaBAD)568 rph-1 ΔnuoK | | kanamycin | Baba *et al*, 2006 (*35*) |
| Keio plate 1 | BW25113 | rrnB DElacZ4787 HsdR 514 DE(araBAD)567 D E(rhaBAD)568 rph-1 | | kanamycin | Baba *et al*, 2006 (*35*) |
| Keio plate 2 | BW25113 | rrnB DElacZ4787 HsdR 514 DE(araBAD)567 D E(rhaBAD)568 rph-1 | | kanamycin | Baba *et al*, 2006 (*35*) |
| BW WT | BW25113 pLumi | E. coli BW25113 (F- , D(araD-araB)567, DlacZ4787(::rrnB-3), λ–, rph-1, D(rhaD-rhaB)568, hsdR514) | pLumi | kanamycin | Lab stock |
| MG1655 donor | | E.coli K-12 MG1655, F- λ- ilvG rfb- 50 rph-1 | | See **Table S6** for antibiotic used for a specific plasmid | Lab stock |

| | | | | | |
|---|---|---|---|---|---|
| MGddap donor | | MG1655 ΔdapA::ErmR | | 2,6-diaminopimelic acid (DAP)<br>See **Table S6** for antibiotic used for a specific plasmid | Dimitrui *et al*, 2024 (*46*) |
| MG1655-nal recipient | | MG1655 Nal$^R$ | | Nalidixic acid | Dimitrui *et al*, 2024 (*46*) |
| MG1655 recipient | | E.coli K-12 MG1655, F- λ- ilvG rfb- 50 rph-1 | | None. | Medaney *et al,* 2016 (*47*)<br>Dimitriui *et al*, 2019 (*48*) |

**Table S3:** Complete list of conditions plotted in Figure 1F.

| **Plasmid** | **Donor strain** | **Recipient strain** | **Donor Antibiotic** | **Dose** | **Recipient Antibiotic** | **Dose** | **Source** |
|---|---|---|---|---|---|---|---|
| pB10 | MGddap | mg1655 | Tet | 10 μg mL$^{-1}$ | 2,6-diaminopimelic acid (DAP) | 0.15 mM | Pulavan *et al*, 2026 (*22*) |
| pB10 | MG1655 | mg1655_nal | Tet | 10 μg mL$^{-1}$ | nalidixic acid | 30 μg mL$^{-1}$ | Pulavan *et al*, 2026 (*22*) |
| pCF12 | MGddap | mg1655 | Amp | 100 μg mL$^{-1}$ | 2,6-diaminopimelic acid (DAP) | 0.15 mM | Pulavan *et al*, 2026 (*22*) |
| pCF12 | MG1656 | mg1655_nal | Amp | 100 μg mL$^{-1}$ | nalidixic acid | 30 μg mL$^{-1}$ | Pulavan *et al*, 2026 (*22*) |
| pCT | MGddap | mg1655 | Amp | 100 μg mL$^{-1}$ | 2,6-diaminopimelic acid (DAP) | 0.15 mM | Pulavan *et al*, 2026 (*22*) |
| pCT | MG1657 | mg1655_nal | Amp | 100 μg mL$^{-1}$ | nalidixic acid | 30 μg mL$^{-1}$ | Pulavan *et al*, 2026 (*22*) |
| pCU1 | MGddap | mg1655 | Strep | 50 μg mL$^{-1}$ | 2,6-diaminopimelic acid (DAP) | 0.15 mM | Pulavan *et al*, 2026 (*22*) |
| pCU1 | MG1658 | mg1655_nal | Strep | 50 μg mL$^{-1}$ | nalidixic acid | 30 μg mL$^{-1}$ | Pulavan *et al*, 2026 (*22*) |
| pESBL283 | MGddap | mg1655 | Amp | 100 μg mL$^{-1}$ | 2,6-diaminopimelic acid (DAP) | 0.15 mM | Pulavan *et al*, 2026 (*22*) |

| | | | | | | | |
|---|---|---|---|---|---|---|---|
| pESBL283 | MG1659 | mg1655_nal | Amp | 100 µg mL$^{-1}$ | nalidixic acid | 30 µg mL$^{-1}$ | Pulavan *et al*, 2026 (*22*) |
| pKAZ3 | MGddap | mg1655 | Tet | 10 µg mL$^{-1}$ | 2,6-diaminopimelic acid (DAP) | 0.15 mM | Pulavan *et al*, 2026 (*22*) |
| pKAZ3 | MG1660 | mg1655_nal | Tet | 10 µg mL$^{-1}$ | nalidixic acid | 30 µg mL$^{-1}$ | Pulavan *et al*, 2026 (*22*) |
| pOXA-48 | MGddap | mg1655 | Amp | 100 µg mL$^{-1}$ | 2,6-diaminopimelic acid (DAP) | 0.15 mM | Pulavan *et al*, 2026 (*22*) |
| pOXA-48 | MG1661 | mg1655_nal | Amp | 100 µg mL$^{-1}$ | nalidixic acid | 30 µg mL$^{-1}$ | Pulavan *et al*, 2026 (*22*) |
| pRK100 | MGddap | mg1655 | Tet | 10 µg mL$^{-1}$ | 2,6-diaminopimelic acid (DAP) | 0.15 mM | Pulavan *et al*, 2026 (*22*) |
| R1 | MG1662 | mg1655 | Kan | 10 µg mL$^{-1}$ | nalidixic acid | 30 µg mL$^{-1}$ | Pulavan *et al*, 2026 (*22*) |
| R388 | MGddap | mg1655 | Trim | 10 µg mL$^{-1}$ | 2,6-diaminopimelic acid (DAP) | 0.15 mM | Pulavan *et al*, 2026 (*22*) |
| R388 | MG1663 | mg1655_nal | Trim | 10 µg mL$^{-1}$ | nalidixic acid | 30 µg mL$^{-1}$ | Pulavan *et al*, 2026 (*22*) |
| R6k | MGddap | mg1655 | Strep | 50 µg mL$^{-1}$ | 2,6-diaminopimelic acid (DAP) | 0.15 mM | Pulavan *et al*, 2026 (*22*) |
| R6k | MG1664 | mg1655_nal | Strep | 50 µg mL$^{-1}$ | nalidixic acid | 30 µg mL$^{-1}$ | Pulavan *et al*, 2026 (*22*) |
| RIP113 | MGddap | mg1655 | Tet | 10 µg mL$^{-1}$ | 2,6-diaminopimelic acid (DAP) | 0.15 mM | Pulavan *et al*, 2026 (*22*) |
| RIP113 | MG1665 | mg1655_nal | Tet | 10 µg mL$^{-1}$ | nalidixic acid | 30 µg mL$^{-1}$ | Pulavan *et al*, 2026 (*22*) |
| wtRP4 | MGddap | mg1655 | Tet | 10 µg mL$^{-1}$ | 2,6-diaminopimelic acid (DAP) | 0.15 mM | Pulavan *et al*, 2026 (*22*) |

| | | | | | | | |
|---|---|---|---|---|---|---|---|
| wtRP4 | MG1666 | mg1655_nal | Tet | 10 μg $mL^{-1}$ | nalidixic acid | 30 μg $mL^{-1}$ | Pulavan *et al*, 2026 (*22*) |
| wtRP4 | BWrifR | BWgentR | Carb | 15 μg $mL^{-1}$ | Gent | 125 μg $mL^{-1}$ | this paper |
| wtRP4 | BWrifR | BWgentR | Carb | 60 μg $mL^{-1}$ | Gent | 125 μg $mL^{-1}$ | this paper |
| wtRP4 | BWrifR | BWgentR | Carb | 120 μg $mL^{-1}$ | Gent | 125 μg $mL^{-1}$ | this paper |
| wtRP4 | BWrifR | BWgentR | Kan | 120 μg $mL^{-1}$ | Gent | 125 μg $mL^{-1}$ | this paper |
| wtRP4 | BWrifR | BWgentR | Tet | 7.5 μg $mL^{-1}$ | Gent | 125 μg $mL^{-1}$ | this paper |
| wtRP4 | BWrifR | BWgentR | Tet | 15 μg $mL^{-1}$ | Gent | 125 μg $mL^{-1}$ | this paper |
| wtRP4 | BWrifR | BWcmR | Carb | 15 μg $mL^{-1}$ | Cm | 30 μg mL-1 | this paper |
| wtRP4 | BWrifR | BWcmR | Carb | 40 μg $mL^{-1}$ | Cm | 30 μg mL-1 | this paper |
| wtRP4 | BWrifR | BWcmR | Carb | 80 μg $mL^{-1}$ | Cm | 30 μg mL-1 | this paper |
| wtRP4 | BWrifR | BWcmR | Carb | 120 μg $mL^{-1}$ | Cm | 30 μg mL-1 | this paper |
| wtRP4 | BWrifR | BWcmR | Kan | 80 μg $mL^{-1}$ | Cm | 30 μg mL-1 | this paper |
| wtRP4 | BWrifR | BWcmR | Kan | 160 μg $mL^{-1}$ | Cm | 30 μg mL-1 | this paper |
| wtRP4 | BWrifR | BWcmR | Kan | 240 μg $mL^{-1}$ | Cm | 30 μg mL-1 | this paper |
| wtRP4 | BWrifR | BWcmR | Tet | 10 μg $mL^{-1}$ | Cm | 30 μg mL-1 | this paper |
| wtRP4 | BWrifR | BWcmR | Tet | 15 μg $mL^{-1}$ | Cm | 30 μg mL-1 | this paper |
| wtRP4 | BWrifR | BWcmR | Tet | 20 μg $mL^{-1}$ | Cm | 30 μg mL-1 | this paper |
| wtRP4 | BWrifR | BWcmR | Tet | 30 μg $mL^{-1}$ | Cm | 30 μg mL-1 | this paper |
| wtRP4 | DA | BWrifR | Carb | 120 μg $mL^{-1}$ | Rif | 40 μg mL-1 | this paper |
| wtRP4 | DA | BWrifR | Kan | 120 μg $mL^{-1}$ | Rif | 40 μg mL-1 | this paper |
| wtRP4 | DA | BWrifR | Tet | 15 μg $mL^{-1}$ | Rif | 40 μg mL-1 | this paper |
| sfRP4 | MG1655-strepR | BWrifR | Carb | 120 μg $mL^{-1}$ | Rif | 40 μg mL-1 | this paper |
| sfRP4 | MG1655-strepR | BWrifR | Kan | 120 μg $mL^{-1}$ | Rif | 40 μg mL-1 | this paper |

**Table S4:** Complete list of conditions plotted in Figure 2D.

| Donor | Recipient | Donor Antibiotic Selection | Recipient Antibiotic Selection |
|---|---|---|---|
| DA-wtRP4 | BWrifR | 15xTet | 6xRif |
| BWrifR-wtRP4 | BWcmR | 7.5xTet | 3xCm |
| DA-wtRP4 | BWrifR | 5xTet | 6xRif |
| DA-wtRP4 | BWrifR | 15xKan | 6xRif |
| DA-wtRP4 | BWrifR | 0xTet | 0xRif |
| DA-wtRP4 | BWrifR | 15xCarb | 6xRif |
| DA-wtRP4 | BWrifR | 15xKan | 6xRif |
| DA-wtRP4 | BWrifR | 6xTet | 5xRif |
| DA-wtRP4 | BWrifR | 6xTet | 15xRif |
| DA-wtRP4 | BWrifR | 6xTet | 0xRif |
| DA-wtRP4 | BWrifR | 5xKan | 0xRif |
| DA-wtRP4 | BWrifR | 15xKan | 0xRif |
| DA-wtRP4 | BWrifR | 0xTet | 5xRif |
| DA-wtRP4 | BWrifR | 0xTet | 10xRif |
| DA-wtRP4 | BWrifR | 0xTet | 15xRif |
| DA-wtRP4 | BWrifR | 5xTet | 0xRif |
| DA-wtRP4 | BWrifR | 10xTet | 0xRif |
| DA-wtRP4 | BWrifR | 15xTet | 0xRif |
| DA-wtRP4 | BWrifR | 5xCarb | 0xRif |
| DA-wtRP4 | BWrifR | 10xCarb | 0xRif |
| DA-wtRP4 | BWrifR | 15xCarb | 0xRif |
| DA-wtRP4 | BWrifR | 2xTet | 6xRif |
| DA-wtRP4 | BWrifR | 10xTet | 6xRif |
| DA-wtRP4 | BWrifR | 30xTet | 6xRif |
| DA-wtRP4 | BWrifR | 6xTet | 2xRif |
| DA-wtRP4 | BWrifR | 6xTet | 10xRif |
| DA-wtRP4 | BWrifR | 6xTet | 30xRif |
| DA-wtRP4 | BWrifR | 2xKan | 0xRif |
| pB10 donor | BW WT | 15xTet | 3xKan |

**Table S5:** Complete list of Keio recipients from plates 1 and 2 used in Fig. 4C Attached as a separate Excel file.

**Table S6:** Complete table of plasmids and their selection doses used.

| Plasmid | Resistances (used for selection*) | Donor strain | Selection dose (ug mL-1) |
|---|---|---|---|
| wtRP4 | tetracycline*, carbenicillin, chloramphenicol, kanamycin | DA838 | 15 Tet |
| sfRP4 | chloramphenicol*, streptamycin, kanamycin | MG1655 (ilvA::placIQ1-tetR-FRT) + sRP4-ptetRA-sfGFP-frt-kn-frt-delta-tetRA-frt-Cm-Frt | 30 Cm |
| pB10 | tetracycline*, streptamycin | BWrifR | 15 Tet |
| pZE31-GFP | chloramphenicol | BW25113 | 30 Cm |
| pZE31-GFP + RP4 | chloramphenicol*, tetracycline* | BWcmR | 30 Cm, 15 Tet |
| pCrimson | gentamicin | BW25113 | 50 Gent |
| pLumi | kanamycin | BW25113 | 25 Kan |
| RP4 | ampicillin, tetracycline, kanamycin, | MG1655 / MGddap | 100Amp, 10 Tet, 50 Kan |
| R1 | ampicillin, chloramphenicol, kanamycin, streptamycin | MG1655 / MGddap | 100 Amp, 20 Cm, 50 Kan, 50 Strep |
| pB10 | tetracycline | MG1655 / MGddap | 10 Tet |
| pCF12 | ampicillin | MG1655 / MGddap | 100 Amp |
| pCT | ampicillin | MG1655 / MGddap | 100 Amp |
| pCU1 | streptamycin | MG1655 / MGddap | 50 Strep |
| pESBL283 | ampicillin | MG1655 / MGddap | 100 Amp |
| pKAZ3 | tetracycline | MG1655 / MGddap | 10 Tet |
| pOXA-48 | ampicillin | MG1655 / MGddap | 100 Amp |
| pRK100 | tetracycline | MG1655 / MGddap | 10 Tet |
| R388 | trimethoprim | MG1655 / MGddap | 10 Trim |
| R6K | streptamycin | MG1655 / MGddap | 50 Strep |
| RIP113 | tetracycline | MG1655 / MGddap | 10 Tet |

**Table S7:** Compete table of donor and recipient combinations used.

| Donor / Recipient Pair | Experiment / Figure |
|---|---|
| BWrifR-wtRP4/BWcmR | Fig.1B, Fig. 1D, Fig.1F, Fig.2D, Fig. SA, Fig.S3C-E, Fig.S4B |
| DA-wtRP4/BWrifR | Fig.1C, Fig.1F, Fig.2B, Fig.2C, Fig.2D, Fig.3D, Fig.4B, Fig.3E, Fig.S4C-D, Fig.S7A, Fig.S7C |
| MG-wtRP4/MG1655_nal | Fig.1E, Fig.1F |
| MG-R1/MG1655_nal | Fig.1E, Fig.1F |
| BWrifR-wtRP4/BWgentR | Fig.1F, Fig.2D, Fig.S4A |

| | |
|---|---|
| MGddap-pB10/MG1655 | Fig.1F |
| MG1655-pB10/MG1655_nal | Fig.1F |
| MGddap-pCF12/MG1655 | Fig.1F |
| MG1655-pCF12/MG1655_nal | Fig.1F |
| MGddap-pCT/MG1655 | Fig.1F |
| MG1655-pCT/MG1655_nal | Fig.1F |
| MGddap-pCU1/MG1655 | Fig.1F |
| MG1655-pCU1/MG1655_nal | Fig.1F |
| MGddap-pESBL283/MG1655 | Fig.1F |
| MG1655-pESBL283/MG1655_nal | Fig.1F |
| MGddap-pKAZ3/MG1655 | Fig.1F |
| MG1655-pKAZ3/MG1655_nal | Fig.1F |
| MGddap-pOXA-48/MG1655 | Fig.1F |
| MG1655-pOXA-48/MG1655_nal | Fig.1F |
| MGddap-pRK100/MG1655 | Fig.1F |
| MG1655-pRK100/MG1655_nal | Fig.1F |
| MGddap-R388/MG1655 | Fig.1F |
| MG1655-R388/MG1655_nal | Fig.1F |
| MGddap-R6K/MG1655 | Fig.1F |
| MG1655-R6K/MG1655_nal | Fig.1F |
| MGddap-RIP113/MG1655 | Fig.1F |
| MG1655-RIP113/MG1655_nal | Fig.1F |
| MGddap-R1/MG1655 | Fig.1F |
| MGddap-RP4/MG1655 | Fig.1F |
| MG-sfRP4/BWrifR | Fig.1F, Fig.S1, Fig.S7A-B |

| | |
|---|---|
| pB10 donor/ 173 Keio recipients (listed in Table S6) | Fig.4C |
| pB10 donor/ Δ*sucA* | Fig.4D, Fig.4F |
| pB10 donor/ Δ*citB* | Fig.4D, Fig.4F |
| pB10 donor/ Δ*fumB* | Fig.4D, Fig.4F |
| pB10 donor/ Δ*nuoK* | Fig.4D, Fig.4F |
| pB10 donor/BW WT | Fig. 2D, Fig.4D, Fig.4F, Fig.S8 |

**Table S8:** Primers used to verify Keio mutants.

| Gene | Primer |
|---|---|
| Keio Δ*sucA* up fwd | AGTATCCACGGCGAAGTAAGCATAAAAAAGATGCTTAAGGGATCACGATG |
| Keio Δ*sucA* down rev | ATCTACGCTACTCATTGTGTATCCTTTATTTATTCGACGTTCAGCGCGTC |
| Keio Δ*citB* up fwd | CCGGTGAACATGGCATTGGG |
| Keio Δ*citB* down rev | TCTTCCGGCATTAACCCGGC |
| Keio Δ*fumB* up fwd | TGGCAAGCAGTAGCTGCACG |
| Keio Δ*fumB* down rev | CCAGAGCTTTATTCTGCCGGG |
| Keio Δ*nuoK* up fwd | GGCAAGCACCAGAACCACC |
| Keio Δ*nuoK* down rev | GAACGCCAGTGGCTGAAACC |
| Kan K2 fwd | CGGTGCCCTGAATGAACTGC |
| Kan K1 rev | CAGTCATAGCCGAATAGCCT |

**Table S9:** Model parameters

| Model component | Parameter | Value | Interpretation | Units | How obtained / where used |
|---|---|---|---|---|---|
| **Growth rate** | mD | 1.37 | Drug-free donor growth rate | h^-1 | Fit to log donor CFU |
| **Initial density** | N0,D | 4354779.28 | Donor abundance at the control-growth fit intercept | CFU | Intercept from donor control |
| **Growth rate** | μR | 1.18 | Drug-free recipient growth rate | h^-1 | Fit to log recipient CFU |
| **Initial density** | N0,R | 7.15E+06 | Recipient abundance at the control-growth fit intercept | CFU | Intercept from recipient control |
| **Antibiotic sensitivity** | KD,Rif | 2.22E+21 | Rif concentration-response coefficient for donor growth | (fold MIC)^-1 | Fit as a near-static donor response to Rif data |
| **Antibiotic sensitivity** | aD,Rif | 0.00 | Linear Rif-dependent donor killing coefficient | h^-1 (fold MIC)^-1 | Set to zero because the Rif donor response was fit with the static K form. |

| | | | | | |
|---|---|---|---|---|---|
| **Antibiotic sensitivity** | KR,Tet | 2354.16 | Tet concentration-response coefficient for recipient growth | (fold MIC)^-1 | Fit as a near-static recipient response to Tet data |
| **Antibiotic sensitivity** | aR,Tet | 0.00 | Linear Tet-dependent recipient killing coefficient | h^-1 (fold MIC)^-1 | Set to zero because the Tet recipient response was fit with the static K form. |
| **Antibiotic sensitivity** | aR,Kan | 1.90 | Linear Kan-dependent recipient killing coefficient | h^-1 (fold MIC)^-1 | Fit on log CFU |
| **Antibiotic sensitivity** | KR,Kan | 0.00 | Kan concentration-response coefficient for recipient growth | (fold MIC)^-1 | Set to zero because Kan was fit with the linear killing form. |
| **Antibiotic sensitivity** | aR,Carb | 0.71 | Linear Carb-dependent recipient killing coefficient | h^-1 (fold MIC)^-1 | Fit on log recipient CFU with a 1 h onset delay |
| **Antibiotic sensitivity** | KR, Carb | 0.00 | Carb concentration-response coefficient for recipient growth | (fold MIC)^-1 | Set to zero because Carb was fit with the linear killing form. |
| **Competence loss** | kD (Rif) | 0.02 | Rate of donor competence loss per Rif fold MIC | h^-1 (fold MIC)^-1 | Jointly fit by log-residual minimization to Tet/Rif background data |
| **Competence loss** | kR (Tet) | 0.04 | Rate of recipient competence loss per Tet fold MIC | h^-1 (fold MIC)^-1 | Jointly fit by log-residual minimization to Tet/Rif background data |
| **Competence loss** | kKan | 0.12 | Rate of recipient competence loss per Kan fold MIC | h^-1 (fold MIC)^-1 | Fit jointly to Kan background data |
| **Competence loss** | kCarb | 3.43E-09 | Rate of recipient competence loss per Carb fold MIC after delay | h^-1 (fold MIC)^-1 | Fit jointly to Carb background data |
| **Conjugation** | h_tube | 1.00E-12 | Tube conjugation coefficient multiplying competent donor and recipient pools | CFU^-1 h^-1 | Obtained from Dimitriu et al |
| **Conjugation** | h_plate | 1.00E-10 | Plate conjugation coefficient | CFU^-1 h^-1 | Calculated as η0 x 100 agar fold increase from Dimitriu et al |
| **Growth rate** | mT | 1.10 | Growth rate of adapted transconjugants at 0× Tet | h^-1 | Exponential log-CFU fit to RP4 Tet/Rif tube data; averaged across all conditions |
| **Initial density** | N0,T | 1.00 | Adapted-transconjugant abundance at fitted intercept | CFU | Assumed initial T density as conservative estimate |
